# Transdiagnostic Symptom Domains are Associated with Head Motion During Multimodal Imaging in Children

**DOI:** 10.1101/2024.09.13.612668

**Authors:** Kavari Hercules, Zhiyuan Liu, Jia Wei, Gladys Venegas, Olivia Ciocca, Alice Dyer, Goeun Lee, Sasha Santini-Bishop, Heather Shappell, Dylan G. Gee, Denis G. Sukhodolsky, Karim Ibrahim

## Abstract

**Background:** Head motion is a challenge for neuroimaging research in developmental populations. However, it is unclear how transdiagnostic symptom domains including attention, disruptive behavior (e.g., externalizing behavior), and internalizing problems are linked to scanner motion in children, particularly across structural and functional MRI. The current study examined whether transdiagnostic domains of attention, disruptive behavior, and internalizing symptoms are associated with scanner motion in children during multimodal imaging.

**Methods:** In a sample of 9,045 children aged 9-10 years in the Adolescent Brain Cognitive Development (ABCD) Study, logistic regression and linear mixed-effects models were used to examine associations between motion and behavior. Motion was indexed using ABCD Study quality control metrics and mean framewise displacement for the following: T1-weighted structural, resting-state fMRI, diffusion MRI, Stop-Signal Task, Monetary Incentive Delay task, and Emotional n-Back task. The Child Behavior Checklist was used as a continuous measure of symptom severity.

**Results:** Greater attention and disruptive behavior problem severity was associated with a lower likelihood of passing motion quality control across several imaging modalities. In contrast, increased internalizing severity was associated with a higher likelihood of passing motion quality control. Increased attention and disruptive behavior problem severity was also associated with increased mean motion, whereas increased internalizing problem severity was associated with decreased mean motion.

**Conclusion:** Transdiagnostic domains emerged as predictors of motion in youths. These findings have implications for advancing development of generalizable and robust brain-based biomarkers, computational approaches for mitigating motion effects, and enhancing accessibility of imaging protocols for children with varying symptom severities.

## Introduction

Movement during functional and structural neuroimaging impacts image quality via the creation of artifacts and reduced data retention (1–4). Successful imaging can be particularly challenging in developmental populations due to increased scanner motion (5, 6). Scanner motion in pediatric populations can also be compounded by symptom domains that make scanning particularly challenging for youths (7). For instance, symptoms associated with neurodevelopmental and other child mental health conditions including attention, externalizing or disruptive behavior (e.g., aggression, irritability/anger, noncompliance), and internalizing problems can make scans challenging and contribute to increased movement (4, 6). Thus, more work is needed to understand the relationships between transdiagnostic symptom domains and motion during multimodal imaging, which will help advance development and validation of safe and effective methods for mitigating head motion while enhancing accessibility of imaging research in youths. In the current study, we examine associations between head motion and transdiagnostic symptom domains of attention, disruptive behavior, and internalizing problems in youths.

Head motion during scanning is a challenge in translational developmental neuroscience research, which can have complex effects on the neural signal depending on features such as the duration, timing, and trajectory of motion (8–11). For instance, head motion often leads to the misalignment of the spatiotemporal units, affecting accurate estimation of the blood oxygenation level dependent (BOLD) signal and potentially obscuring or impacting neural correlates of structure and function (5, 8, 12, 13). Despite post-acquisition approaches for addressing motion—including motion correction via movement parameters entered as covariates in the general linear model (13), global signal regression (14, 15), censoring or scrubbing approaches (8), and denoising (10, 16–19)—motion-related confounds remain a concern as well as lower data retention and impact accessibility of fMRI research for pediatric populations, thus hindering development of robust and reliable brain-based biomarkers (12, 20, 21).

Effects of motion artifact can substantially affect estimates of and/or introduce spurious differences in functional connectivity, particularly between groups of interest such as clinical vs unaffected samples (5, 8, 22). For instance, movement has potential to obscure true effects and/or artificially inflate between-group differences in functional connectivity (6, 8, 10, 12, 23, 24). Head motion has also been shown to affect estimates of functional connectivity and could be misinterpreted as neuronal effects, particularly in cognitive control and social cognitive circuitry including the frontoparietal and default mode networks (22). Along these lines, head motion can impact estimates of functional connectivity, particularly related to resting-state functional MRI (fMRI) (5, 8, 22, 25), and tends to increase connectivity among regions although this may vary depending on the distance between nodes (5, 18, 26). It is also possible that between-group differences in cognitive processes and neural markers attributed to age may be exaggerated due to differences in motion between children of different ages (1, 5, 11, 20, 24, 27). For instance, head motion demonstrates a U-shaped curve across the lifespan such that higher motion is observed at both younger and older ages (20). Thus, motion effects on functional connectivity measures may be particularly pronounced when comparing children vs adults in individual imaging studies or in meta-analyses that often test age effects of cognitive constructs (5, 22, 24, 28).

Clinical subgroups in pediatric populations may be more prone to motion during scanning and data exclusion due to co-occurring symptoms that can make fMRI challenging, particularly attention, disruptive behavior and internalizing problems. Relatively fewer studies have examined motion in clinical subgroups vs unaffected controls in children compared to community-based and/or adult samples. The few existing studies have shown lower scan success rates and greater motion in clinical samples including children with autism spectrum disorder (ASD), attention-deficit/hyperactivity disorder (ADHD), and epilepsy than unaffected controls (4, 7, 29). Studies have also shown associations between greater motion and attention-related ADHD symptoms in youths (7, 30, 31). However, no studies to date have examined how motion is related to transdiagnostic domains of symptoms in youths, particularly disruptive behavior and internalizing problems.

Despite advances in preprocessing, motion correction, and denoising methodologies, the extent to which head motion in youths impacts neural activation and connectivity estimates remains relatively unknown considering the variability across study findings (1, 5, 12). While exclusion of high-motion participants mediates motion artifacts for connectivity estimates, this may introduce a selection bias by systematically altering the study population. For instance, low-motion clinical groups may be more phenotypically similar to unaffected, typically developing controls compared to excluded high-motion participants, thereby reducing potential between-group differences (4). Prior studies of pediatric scan motion were also limited by sample sizes, lack of focus on transdiagnostic symptom domains, and/or limited range of imaging modalities because the majority of studies have focused on task-based or resting-state fMRI (21). Given that motion may disproportionately affect clinical developmental populations (7), this context bolsters the need for increased understanding of motion and its predictors to develop well-informed protocols for scanning in youths. Identifying predictors of scanner motion in pediatric samples and the link to transdiagnostic symptom domains has the potential to inform future studies and efforts to develop methods for maximizing data retention and enhancing accessibility of fMRI studies for children.

The current study examined associations between transdiagnostic symptom domains of attention, disruptive behavior, and internalizing problems and head motion during functional and structural neuroimaging modalities in youths from the Adolescent Brain Cognitive Development (ABCD) Study℠ (32, 33). First, we tested if transdiagnostic symptom domains were associated with passing scan quality control across imaging modalities (T1-weighted structural, resting-state fMRI, task-based fMRI, and diffusion MRI) as determined by the ABCD inclusion flag variables (dichotomously coded as 1 = pass or 0 = fail imaging quality control) (33). The task-based fMRI scans included the Stop-Signal Task (SST) that measures inhibitory control, the Monetary Incentive Delay (MID) task that measures reward-based learning, and the Emotional N-back (EN-Back) task that measures executive control and emotion perception (32, 33). For a continuous measure of symptom severities, we used the Child Behavior Checklist (CBCL) (34) broadband scales (Attention, Externalizing, and Internalizing Problems scales). Next, in the sample of participants passing motion quality control, we examined if transdiagnostic symptom domains were associated with mean motion modeled as a continuous variable for each modality. As a follow-up supplemental analysis, we also tested for the moderating role of sex in all models. Based on previous work (1, 2, 4, 7, 22, 29), we expected greater severity of symptom domains to be associated with a reduced likelihood of passing motion quality control across structural and functional MRI. We also hypothesized that associations would emerge between greater severity of symptom domains and greater mean head motion during multimodal imaging. Given that no prior work, to our knowledge, has directly examined associations between internalizing severity and head motion, we did not have a priori directional hypotheses for this symptom domain. Nonetheless, we expected internalizing severity to be associated with head motion. Exploratorily, we also examined associations between motion and demographic variables such as age, sex, and cognitive performance for comparison to prior work.

## Materials and Methods

### Participants

We analyzed a subset of participants from the ABCD Study® (32) (https://abcdstudy.org/; baseline study measures from ABCD [release 4.0]). The ABCD Study is a multi-site longitudinal study with over 11,500 children aged 9-10 years during the first wave of enrollment, that comprehensively characterizes cognitive and neural development from early adolescence to early adulthood (32). At baseline, there were approximately 11,879 subjects, and 2,834 were removed due to missing and incomplete records, resulting in 9,045 children (4,408 females) with behavioral and imaging data available for analysis. First, using logistic regression, we utilized the sample of 9,045 subjects to predict the probability of passing vs failing quality control for each scanning imaging modality. Next, using linear mixed-effects models, we utilized the sample of youths who passed quality control for each imaging modality for analyses with continuous measures of symptom severity (n=8,677 for T1-weighted, n=7,719 for resting-state, n=7,431 for diffusion MRI, n=7,115 for SST task, n=7,626 for MID task, n=6,875 for EN-back task) (**Figure 1**). Based on ABCD imaging analysis recommendations (33), the provided inclusion variables or flags were used for analyses in which participants passed or failed protocol compliance and quality control for T1-weighted, diffusion MRI, task-based fMRI, and resting-state fMRI scans. Participant characteristics are shown in **Table 1**.

**Figure 1.**
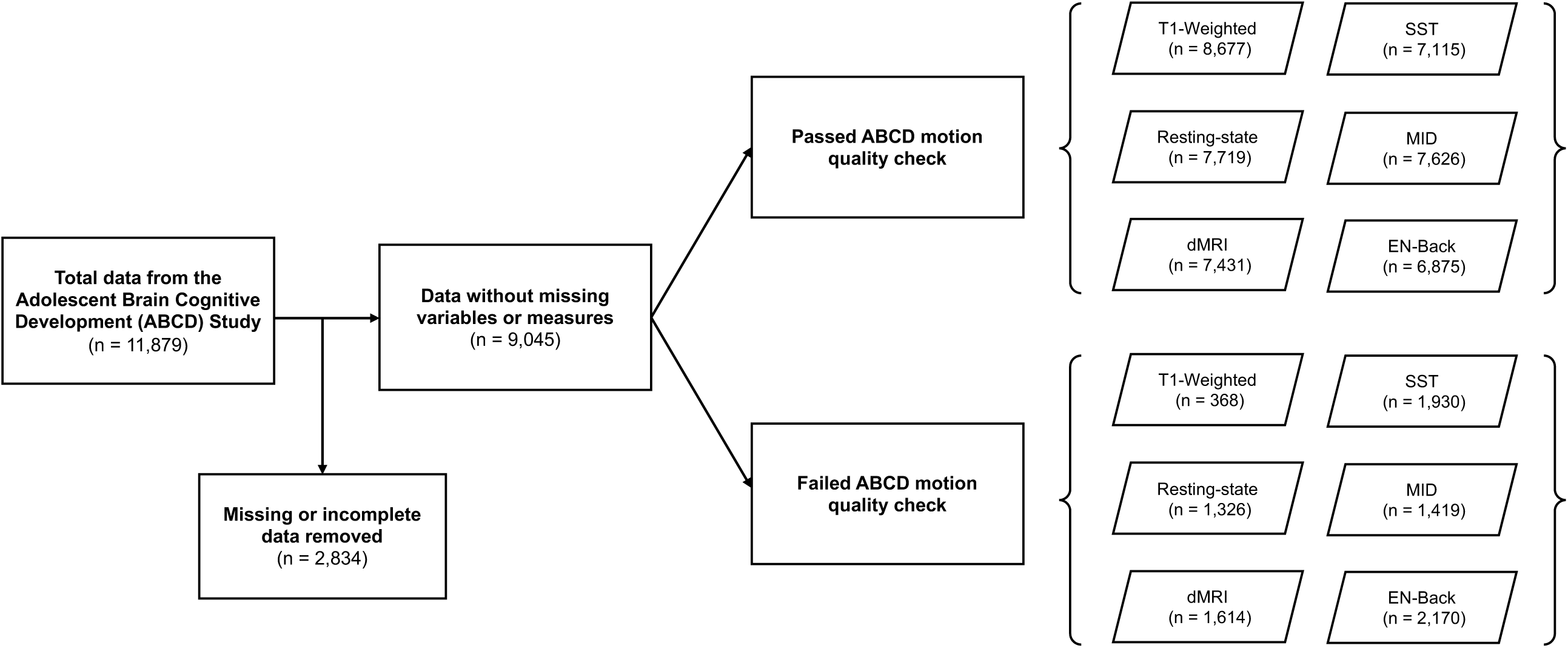
Flowchart illustrating data structure and participants from the ABCD Study dataset. Analyses were conducted for each of the imaging modalities: resting-state fMRI (rsfMRI), diffusion MRI (dMRI), Stop Signal Task (SST), Monetary Incentive Delay (MID) task, and the emotional version of the n-back task (EN-Back).

**Table 1.** Participant Demographics and Characteristics.

| Variable | Total | T-1 Weighted |  |  | Resting-state fMRI |  |  | Diffusion MRI |  |  | SST |  |  | MID |  |  | EN-Back |  |  |
| --- | --- | --- | --- | --- | --- | --- | --- | --- | --- | --- | --- | --- | --- | --- | --- | --- | --- | --- | --- |
|  |  | Pass QC | Fail QC | *p* value<sup>b</sup> | Pass QC | Fail QC | *p* value<sup>b</sup> | Pass QC | Fail QC | *p* value<sup>b</sup> | Pass QC | Fail QC | *p* value<sup>b</sup> | Pass QC | Fail QC | *p* value<sup>b</sup> | Pass QC | Fail QC | *p* value<sup>b</sup> |
|  | 9,045 | 8,677 | 368 |  | 7,719 | 1,326 |  | 7,431 | 1,614 |  | 7,115 | 1,930 |  | 7,626 | 1,419 |  | 6,875 | 2,170 |  |
| Age, years (SD) | 9.9 (7.5) | 9.9 (7.5) | 9.8 (7.3) | 0.001 | 9.9 (7.5) | 9.8 (7.2) | 1.27e-13 | 9.9 (7.5) | 9.9 (7.4) | 0.043 | 9.9 (7.5) | 9.9 (7.4) | 1.57e-08 | 9.9 (7.5) | 9.9 (7.4) | 0.0654 | 10.0 (7.5) | 9.8 (7.3) | 4.18e-23 |
| Male, % | 51.3 | 51.1 | 54.6 | 0.207 | 49.6 | 61.1 | 1.21e-14 | 50.9 | 52.9 | 0.168 | 50.8 | 53.1 | 0.0802 | 50.2 | 56.8 | 6.37e-06 | 51.3 | 51.1 | 0.884 |
| Race, % |  |  |  |  |  |  |  |  |  |  |  |  |  |  |  |  |  |  |  |
| White | 54.2 | 54.5 | 46.5 | 0.003 | 55.1 | 48.9 | 4e-05 | 54.0 | 55.1 | 0.436 | 56.4 | 45.8 | 0.436 | 54.8 | 50.7 | 0.00514 | 57.8 | 42.7 | 8.16e-35 |
| Black | 13.1 | 13.0 | 16.6 | 0.055 | 12.5 | 16.6 | 6.6e-05 | 13.6 | 11.2 | 0.011 | 11.6 | 18.8 | 0.0105 | 12.4 | 16.8 | 8.14e-06 | 10.7 | 20.7 | 3.99e-33 |
| Asian | 2.1 | 2.1 | 3.0 | 0.312 | 2.0 | 2.6 | 0.179 | 1.9 | 2.9 | 0.018 | 2.2 | 1.9 | 0.0177 | 2.0 | 2.6 | 0.189 | 2.2 | 1.8 | 0.362 |
| Other | 10.4 | 10.2 | 14.1 | 0.019 | 10.2 | 11.4 | 0.195 | 10.4 | 10.2 | 0.82 | 9.8 | 12.3 | 0.82 | 10.1 | 11.7 | 0.0766 | 10.1 | 11.2 | 0.126 |
| Ethnicity, % |  |  |  |  |  |  |  |  |  |  |  |  |  |  |  |  |  |  |  |
| Hispanic | 20.2 | 20.3 | 19.8 | 0.899 | 20.2 | 20.4 | 0.869 | 20.1 | 20.7 | 0.635 | 20.0 | 21.2 | 0.635 | 20.6 | 18.1 | 0.0332 | 19.2 | 23.5 | 1.57e-05 |
| Cognition<sup>a</sup>, mean (SD) | 101.4 (17.5) | 101.5 (17.5) | 98.5 (17.7) | 0.001 | 102.1 (17.5) | 97.6 (17.6) | 3.55e-17 | 101.3 (17.4) | 102.0 (18.3) | 0.146 | 102.7 (17.3) | 96.6 (17.6) | 1.96e-40 | 101.8 (17.4) | 99.2 (18.4) | 5.18e-07 | 103.8 (16.9) | 93.8 (17.5) | 1.12e-112 |
| CBCL Attention, mean (SD) | 2.8 (3.4) | 2.8 (3.4) | 3.4 (3.6) | 0.003 | 2.7 (3.3) | 3.5 (3.7) | 3.19e-14 | 2.8 (3.4) | 2.8 (3.3) | 0.541 | 2.7 (3.3) | 3.5 (3.8) | 1.13e-16 | 2.8 (3.4) | 3.2 (3.5) | 0.0001 | 2.7 (3.3) | 3.4 (3.7) | 7.38e-17 |
| CBCL Externalizing, mean (SD) | 4.3 (5.7) | 4.3 (5.7) | 4.3 (5.5) | 0.947 | 4.1 (5.5) | 5.0 (6.5) | 3.25e-06 | 4.3 (5.7) | 4.0 (5.3) | 0.038 | 4.0 (5.4) | 5.1 (6.5) | 2.48e-10 | 4.2 (5.6) | 4.6 (6.1) | 0.009 | 4.0 (5.4) | 5.0 (6.4) | 3.63e-11 |
| CBCL Internalizing, mean (SD) | 5.0 (5.5) | 5.0 (5.5) | 4.8 (5.5) | 0.526 | 4.9 (5.4) | 5.3 (5.8) | 0.0285 | 5.0 (5.5) | 4.7 (5.3) | 0.027 | 4.9 (5.4) | 5.2 (5.8) | 0.0619 | 4.9 (5.4) | 5.1 (5.6) | 0.292 | 4.9 (5.3) | 5.2 (5.9) | 0.0498 |
| FD, mean (SD) |  |  |  |  | 0.3 (0.24) | 0.9 (0.79) | 2.57e-133 | 1.3 (0.48) | 1.6 (0.88) | 1e-44 | 0.4 (0.40) | 0.7 (0.95) | 3.92e-42 | 0.4 (0.36) | 0.6 (0.82) | 2.08e-20 | 0.4 (0.4) | 0.8 (0.95) | 3.58e-53 |

Note: CBCL, Child Behavior Checklist; EN-Back, Emotional N-Back Task; MID, Monetary Incentive Delay Task; SST, Stop Signal Task; FD, Framewise Displacement; QC, Quality Control (based on the Adolescent Brain Cognitive Development study protocol procedures (32, 33)).
General Cognition measured by the NIH Toolbox (35).
Significant group differences at $p < 0.05$ using Chi-square test for categorical variables or independent samples T-test.

### Behavioral Measures

We used a broadband continuous measure of symptoms related to child mental health, which was selected to allow comparison with prior imaging studies, particularly using the ABCD dataset. The parent-rated Child Behavior Checklist (CBCL) scores (34) were used as a transdiagnostic, dimensional measure in analyses for severity of attention, disruptive behavior, and internalizing problems. The CBCL is a well-established measure of child psychopathology. The *CBCL Externalizing Problems* scale includes items reflecting disruptive behaviors related to verbal and physical aggression, conduct problems (e.g., attacks, setting fires, running away, rule-breaking, truancy), anger/irritability, and noncompliance. The *CBCL Internalizing Problems* scale includes items related to somatic complaints, social withdrawal, and anxiety/depressed symptoms. The *CBCL Attention Problems* scale includes symptoms related to inattention, hyperactivity, and impulsivity. Participants also completed cognitive assessments including the NIH Toolbox Cognition Battery (Picture Vocabulary, Flanker Test, List Sort Working Memory Task, Dimensional Change Card Sort Task, Pattern Comparison Processing Speed Task, Picture Sequence Memory Task, and the Oral Reading Test) (35). The NIH Toolbox Cognition Battery age-corrected total composite score was used in analyses to account for overall cognitive performance.

### ABCD Study: Imaging data, processing, and task descriptions

The design and imaging protocol of the larger ABCD Study has been described in prior work (32, 33). Details of ABCD Study recruitment (36), neurocognitive batteries (37), and imaging protocols (33) are also available elsewhere. The released imaging data (T1-weighted, diffusion weighted imaging, resting-state, and task-based fMRI) were processed through ABCD’s Data Analysis, Informatics and Resource Center (DAIRC) image processing pipeline (33). The ABCD scan session includes a structural T1-weighted series (33), diffusion MRI series using a multiband EPI (33, 38, 39) that includes 96 diffusion directions (6 directions at b=500 s/mm^2^, 15 at b=1000 s/mm^2^, 15 at b=2000 s/mm^2^, and 60 at b=3000 s/mm^2^), a resting-state series that includes at least 8 minutes of viewing a blank screen with a cross-hair with eyes open, and a task-based fMRI battery (33). The ABCD task-based fMRI battery includes the Stop-Signal Task (SST), Emotional N-back task (EN-Back), and Monetary Incentive Delay (MID) task (32). The SST is designed to engage circuitry of inhibition and error monitoring and requires rapid responding to a “Go” stimulus, unless followed by a second “Stop” stimulus prompting participants to cancel their response. The EN-Back task is designed to engage circuitry of working memory, with a value-added probe of social information processing involving blocks of images of places vs emotional or neutral faces. The MID task is designed to engage reward circuitry and requires anticipating and experiencing different magnitudes of reward and loss. Together, these tasks tap cognitive processes and neural circuitry related to emotion perception, response inhibition, reward anticipation, and cognitive control more broadly including error processing and working memory (32, 40–44), and were selected based on processes implicated in risk of substance use and child psychopathology (45, 46). For more detail regarding the ABCD Study scan sequences and acquisition parameters, see Casey et al. (32) and Hagler et al. (33).

Within the ABCD data release, a recommended sample for each imaging modality is provided by the DAIRC using a dichotomously coded variable. Thus, for each imaging series, we used the provided binary variables indicating if participants had adequate data (e.g., >375 frames, passed FreeSurfer quality control) and high quality data or passing ABCD quality control (variables “imgincl_{t1w, rsfmri, dmri, mid, nback, sst}_include” of the “abcd_imgincl01” instrument). Additional detail is provided in the **Supplemental Methods** and **Table S1**. The DAIRC also provides values for mean framewise displacement (FD) (8), a commonly used metric of head motion, which was used for linear mixed-effects analyses predicting motion as a continuous variable. FD represents the sum of movement across the six rigid body motion parameters and across a scan series.

### Statistical Analysis

First, the normality of behavioral variables was assessed based on the degree of skewness (using ‘skewness’ from the ‘e1071’ R package). The CBCL subscale scores (Attention, Externalizing, and Internalizing Problems scales) were non-normally distributed (i.e., skewness score >1) and log transformed (using ‘optLogTransform’ from the ‘optLog’ R package). As a lower absolute value in skewness indicates better normality, after log transformation, skewness for Attention Problems was reduced from 1.44 to -0.0026, Internalizing Problems was reduced from 1.94 to 0.00042, and Externalizing Problems was reduced from 2.3 to -0.014. Other variables for age and cognitive performance were normally distributed (i.e., all skewness scores <0.3). Next, missing data was imputed from the mean for cognitive performance scores from NIH Toolbox (<1% of records). Lastly, results of pair-wise Pearson correlations among variables showed values <0.59 and non-zero eigenvalues (ranging from 0.38-2.1) along with the Variance Inflation Factor (VIF) values <1, all of which indicate low risk of multicollinearity (47) (**Figure 2**).

**Figure 2.**
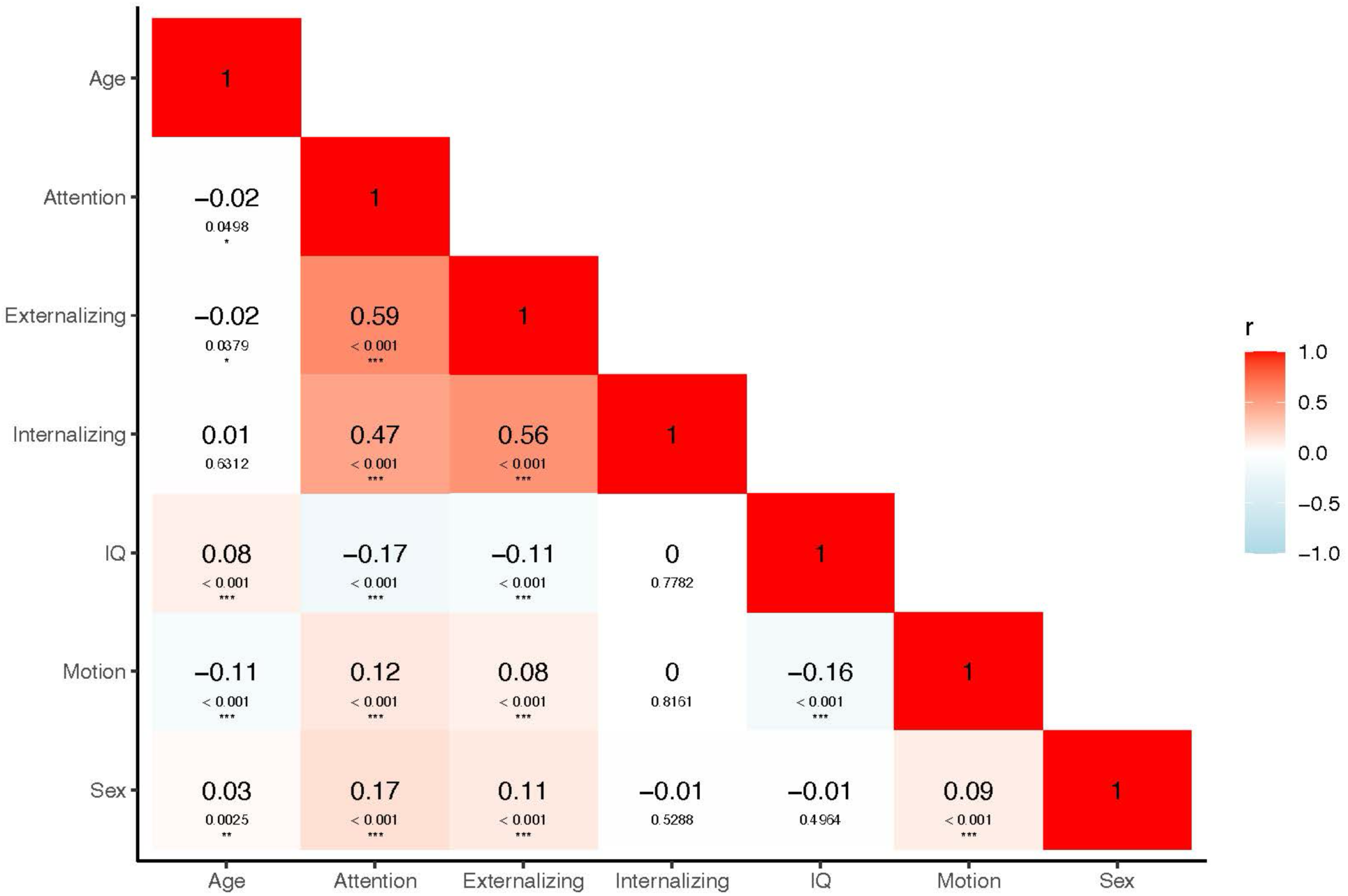
Correlations among study variables. Pearson correlations and significance values are shown for age, CBCL scales (Attention, Externalizing, and Internalizing Problems scores), cognition (IQ), motion indexed by mean framewise displacement (mm), and sex at birth. The matrix values indicate the correlation coefficient (top) and the corresponding p-values (bottom). The strength and direction of the correlations are color-coded, with red representing positive correlations and blue representing negative correlations. Statistically significant correlations are denoted by asterisks: ***p < 0.001, **p < 0.01, and *p < 0.05.

#### Logistic Regression Models

In the total sample (*n*=9,045), we applied logistic regression models in R (via ‘glmer’) to test if transdiagnostic domains of psychopathology (CBCL Attention, Externalizing, and Internalizing Problems scales) are associated with passing (n ≈ 7,300) vs failure (n ≈ 1,745) of scan quality control, modeled as a binary dependent variable (1=pass, 0=fail) based on ABCD DAIRC inclusion variables for each imaging modality (**Figure 1**). All models included the following independent variables: age, sex at birth, race/ethnicity, cognitive performance, and symptom domains (CBCL Attention, Externalizing, and Internalizing Problems scores). Random intercepts were included for study site and family (i.e., having a sibling in the study) nested within the site. Follow-up, supplemental analyses were conducted to test for potential effects of sex. Logistic regression models were repeated with an interaction term for sex with each behavioral domain (i.e., Sex-by-CBCL Attention Problems, Sex-by-CBCL Externalizing Problems, and Sex-by-CBCL Internalizing Problems). We fitted logistic regression models for each of the six imaging modalities: T1-weighted, resting-state fMRI, diffusion MRI, and task fMRI (SST, MID, and EN-back tasks). All results and final p-values were then FDR-corrected across all tests. To facilitate interpretation of results and effect sizes, Odds Ratios were calculated for each variable across each scanning series or imaging modality.

#### Linear Mixed-Effects Models

In the sample of participants who passed ABCD quality control, we then applied linear mixed-effects models in R (via ‘lmer’) to test the association between transdiagnostic domains of behavior and motion modeled as a continuous variable using mean FD as the dependent variable. All models included the following independent variables: age, sex at birth, race/ethnicity, cognitive performance, and behavioral domains (CBCL Attention, Externalizing, and Internalizing Problems scores). Random intercepts were included for study site and family (i.e., having a sibling in the study) nested within the site. Follow-up, supplemental analyses were also conducted to test for potential effects of sex. Linear mixed-effects models were repeated as above with an interaction term for sex with each behavioral domain (i.e., Sex-by-CBCL Attention Problems, Sex-by-CBCL Externalizing Problems, and Sex-by-CBCL Internalizing Problems). We fitted linear mixed-effects models for each of the five relevant imaging modalities with a mean FD metric: resting-state fMRI, diffusion MRI, and task fMRI (SST, MID, and EN-back tasks). All results and final p-values were FDR-corrected across all tests. To facilitate interpretation of results and effect sizes, we also converted the coefficients for the linear mixed-effects models to Cohen’s d.

### Data and Code Availability

Data from the ABCD Study are shared on the National Institute of Mental Health Data Archive (NDA) https://nda.nih.gov/. To promote transparency, all code used for analyses is available on GitHub (https://github.com/EmotionNeuroscienceLab).

## Results

### Transdiagnostic Domains of Behavior and Imaging Quality Control

Regarding attention problems, significant associations were identified with passing motion quality control across all imaging modalities (**Table 2**). That is, individuals with increased severity of attention problems were less likely to pass motion quality control for each imaging modality. Specific modalities showed a more pronounced decrease in odds, indicating a lower likelihood of passing the motion quality checks (**Table 2**), and these included the T1-weighted scan (25.8% decrease, *p_FDR_* = 0.0002), resting-state fMRI (20.2% decrease, *p_FDR_* = 4.65e-07), diffusion MRI (19.9% decrease, *p* = 0.0005), and the SST task (15.4% decrease, *p_FDR_* = 1.76e-05). Effects for the MID and EN-Back tasks were less pronounced with 10.1% (*p_FDR_* = 0.04) and 10.8% (*p_FDR_* = 0.01) decreases in odds, respectively. Regarding internalizing and disruptive behavior problems, significant associations were observed for the SST and EN-Back tasks (all *ps* <0.05). That is, increased internalizing problem severity was linked with a 10% increase (*p* = 0.01) in the likelihood of passing motion quality control during the SST task, while increased disruptive behavior problem severity was linked with a 9.4% decrease (*p_FDR_* = 0.01) in the likelihood of passing motion quality control during the EN-Back task.

**Table 2.** Results of Logistic Regression Models Predicting Scan Quality Control and Domains of Transdiagnostic Symptoms.

| Variable | Estimate | Std. Error | z value | p | p<sub>FDR</sub> | OR | 95% CI |  |
| --- | --- | --- | --- | --- | --- | --- | --- | --- |
|  |  |  |  |  |  |  | 2.5% | 97.5% |
| T1-Weighted |  |  |  |  |  |  |  |  |
| Intercept | 1.68 | 0.940 | 1.79 | 0.0731 | 0.156 | 5.39 | 0.85 | 34 |
| Age | 0.012 | 0.007 | 1.64 | 0.1 | 0.189 | 1.01 | 0.99 | 1.03 |
| Sex | -0.097 | 0.110 | -0.89 | 0.377 | 0.508 | 0.91 | 0.73 | 1.13 |
| Black | -0.210 | 0.171 | -1.23 | 0.219 | 0.345 | 0.81 | 0.58 | 1.13 |
| Asian | 0.075 | 0.163 | 0.46 | 0.643 | 0.693 | 1.08 | 0.78 | 1.48 |
| Other | -0.378 | 0.332 | -1.14 | 0.254 | 0.381 | 0.69 | 0.36 | 1.31 |
| Hispanic | -0.096 | 0.169 | -0.57 | 0.572 | 0.664 | 0.91 | 0.65 | 1.27 |
| Cognition | 0.007 | 0.003 | 2.05 | 0.0403 | 0.0986 | 1.01 | 1 | 1.01 |
| CBCL Attention | -0.298 | 0.073 | -4.11 | 4e-05 | **0.0002** | 0.74 | 0.64 | 0.89 |
| CBCL Externalizing | 0.135 | 0.072 | 1.89 | 0.0587 | 0.129 | 1.15 | 0.99 | 1.32 |
| CBCL Internalizing | 0.114 | 0.067 | 1.70 | 0.0899 | 0.175 | 1.12 | 0.98 | 1.28 |
| Resting-state fMRI |  |  |  |  |  |  |  |  |
| Intercept | -2.240 | 0.547 | -4.1 | 4.11e-05 | 0.000208 | 0.11 | 0.04 | 0.31 |
| Age | 0.027 | 0.004 | 6.34 | 2.3e-10 | 2.53e-09 | 1.03 | 1.02 | 1.04 |
| Sex | -0.448 | 0.065 | -6.95 | 3.74e-12 | 4.94e-11 | 0.64 | 0.56 | 0.73 |
| Black | -0.066 | 0.101 | -0.67 | 0.512 | 0.617 | 0.934 | 0.77 | 1.14 |
| Asian | -0.077 | 0.096 | -0.80 | 0.422 | 0.557 | 0.93 | 0.77 | 1.12 |
| Other | -0.430 | 0.206 | -2.09 | 0.0369 | 0.094 | 0.65 | 0.43 | 0.97 |
| Hispanic | -0.078 | 0.105 | -0.75 | 0.456 | 0.579 | 0.93 | 0.75 | 1.14 |
| Cognition | 0.012 | 0.002 | 6.02 | 1.75e-09 | 1.65e-08 | 1.01 | 1.01 | 1.02 |
| CBCL Attention | -0.226 | 0.042 | -5.43 | 5.64e-08 | **4.65e-07** | 0.79 | 0.74 | 0.8 |
| CBCL Externalizing | 0.013 | 0.042 | 0.31 | 0.752 | 0.788 | 1.01 | 0.93 | 1.1 |
| CBCL Internalizing | 0.025 | 0.039 | 0.65 | 0.514 | 0.617 | 1.02 | 0.95 | 1.11 |
| Diffusion MRI |  |  |  |  |  |  |  |  |
| Intercept | 0.207 | 1.06 | 0.19 | 0.845 | 0.858 | 1.23 | 0.15 | 9.82 |
| Age | 0.008 | 0.006 | 1.45 | 0.148 | 0.254 | 1.01 | 0.99 | 1.02 |
| Sex | -0.114 | 0.089 | -1.28 | 0.201 | 0.331 | 0.89 | 0.75 | 1.06 |
| Black | -0.023 | 0.143 | -0.16 | 0.873 | 0.873 | 0.98 | 0.74 | 1.29 |
| Asian | 0.15 | 0.131 | 1.14 | 0.253 | 0.381 | 1.16 | 0.89 | 1.5 |
| Other | -0.242 | 0.269 | -0.89 | 0.37 | 0.508 | 0.76 | 0.46 | 1.33 |
| Hispanic | -0.061 | 0.136 | -0.45 | 0.651 | 0.693 | 0.94 | 0.72 | 1.23 |
| Cognition | 0.004 | 0.003 | 1.44 | 0.15 | 0.254 | 1 | 0.99 | 1.01 |
| CBCL Attention | -0.222 | 0.058 | -3.82 | 0.000132 | **0.0005** | 0.80 | 0.71 | 0.89 |
| CBCL Externalizing | 0.073 | 0.058 | 1.25 | 0.213 | 0.343 | 1.08 | 0.96 | 1.21 |
| CBCL Internalizing | 0.043 | 0.054 | 0.79 | 0.431 | 0.558 | 1.04 | 0.94 | 1.16 |
| **SST task** |  |  |  |  |  |  |  |  |
| Intercept | -1.85 | 0.463 | -3.99 | 6.52e-05 | 0.000307 | 0.16 | 0.06 | 0.39 |
| Age | 0.014 | 0.004 | 3.9 | 9.7e-05 | 0.000427 | 1.01 | 1.01 | 1.02 |
| Sex | -0.036 | 0.054 | -0.67 | 0.505 | 0.617 | 0.97 | 0.87 | 1.07 |
| Black | -0.243 | 0.085 | -2.87 | 0.0041 | 0.0143 | 0.78 | 0.66 | 0.93 |
| Asian | -0.039 | 0.081 | -0.49 | 0.627 | 0.693 | 0.962 | 0.82 | 1.13 |
| Other | -0.098 | 0.196 | -0.50 | 0.617 | 0.693 | 0.907 | 0.62 | 1.33 |
| Hispanic | -0.217 | 0.089 | -2.45 | 0.0141 | 0.0424 | 0.805 | 0.67 | 0.96 |
| Cognition | 0.016 | 0.002 | 9.21 | 3.14e-20 | 6.92e-19 | 1.02 | 1.01 | 1.02 |
| CBCL Attention | -0.167 | 0.035 | -4.72 | 2.39e-06 | **1.76e-05** | 0.846 | 0.78 | 0.91 |
| CBCL Externalizing | -0.077 | 0.036 | -2.11 | 0.0349 | 0.0922 | 0.926 | 0.86 | 0.99 |
| CBCL Internalizing | 0.092 | 0.033 | 2.77 | 0.0056 | **0.0176** | 1.1 | 1.03 | 1.17 |
| **MID task** |  |  |  |  |  |  |  |  |
| Intercept | 0.763 | 0.51 | 1.49 | 0.135 | 0.241 | 2.15 | 0.79 | 5.83 |
| Age | 0.004 | 0.004 | 0.88 | 0.377 | 0.508 | 1 | 0.99 | 1.01 |
| Sex | -0.25 | 0.061 | -4.11 | 3.94e-05 | 0.000208 | 0.78 | 0.69 | 0.88 |
| Black | -0.164 | 0.095 | -1.72 | 0.0855 | 0.171 | 0.85 | 0.70 | 1.02 |
| Asian | 0.186 | 0.092 | 2.02 | 0.0432 | 0.102 | 1.2 | 1.01 | 1.44 |
| Other | -0.317 | 0.197 | -1.6 | 0.109 | 0.199 | 0.73 | 0.49 | 1.07 |
| Hispanic | -0.056 | 0.099 | -0.56 | 0.574 | 0.664 | 0.95 | 0.78 | 1.15 |
| Cognition | 0.007 | 0.002 | 3.66 | 0.000252 | 0.000977 | 1.01 | 1.0 | 1.01 |
| CBCL Attention | -0.095 | 0.039 | -2.42 | 0.0153 | **0.044** | 0.91 | 0.84 | 0.98 |
| CBCL Externalizing | -0.011 | 0.040 | -0.28 | 0.782 | 0.807 | 0.99 | 0.91 | 1.07 |
| CBCL Internalizing | 0.017 | 0.037 | 0.46 | 0.643 | 0.693 | 1.02 | 0.95 | 1.09 |
| **EN-Back task** |  |  |  |  |  |  |  |  |
| Intercept | -5.15 | 0.466 | -11.1 | 2.15e-28 | 7.1e-27 | 0.01 | 0.002 | 0.014 |
| Age | 0.027 | 0.004 | 7.67 | 1.72e-14 | 2.83e-13 | 1.03 | 1.02 | 1.04 |
| Sex | 0.049 | 0.053 | 0.92 | 0.356 | 0.508 | 1.05 | 0.95 | 1.17 |
| Black | -0.348 | 0.082 | -4.23 | 2.34e-05 | 0.000155 | 0.71 | 0.60 | 0.83 |
| Asian | -0.167 | 0.078 | -2.15 | 0.0312 | 0.0858 | 0.85 | 0.73 | 0.99 |
| Other | -0.196 | 0.194 | -1.01 | 0.312 | 0.457 | 0.82 | 0.57 | 1.2 |
| Hispanic | -0.157 | 0.089 | -1.76 | 0.0778 | 0.161 | 0.86 | 0.72 | 1.02 |
| Cognition | 0.032 | 0.002 | 17.2 | 2.59e-66 | 1.71e-64 | 1.03 | 1.03 | 1.04 |
| CBCL Attention | -0.103 | 0.035 | -2.99 | 0.00278 | **0.0102** | 0.90 | 0.84 | 0.97 |
| CBCL Externalizing | -0.099 | 0.036 | -2.78 | 0.00546 | **0.0176** | 0.91 | 0.84 | 0.97 |
| CBCL Internalizing | 0.064 | 0.033 | 1.95 | 0.0513 | 0.117 | 1.07 | 1.0 | 1.14 |

Note: CBCL, Child Behavior Checklist; EN-Back, Emotional N-Back Task; MID, Monetary Incentive Delay Task; OR, odds ratio; SST, Stop Signal Task; General Cognition is measured by the NIH Toolbox age corrected scores (36). Odds ratios are reported relative to the reference group in the case of categorical variables. Odds ratios > 1 indicate increased likelihood of passing quality control among the nonreference group relative to the reference group (i.e., lower likelihood of passing quality control in the reference group). While odds ratios < 1 indicate decreased likelihood of passing quality control. For continuous measures, the odds ratios represent the change in odds per unit change in the measure.

#### Interactions with Sex and Symptom Domains

Analyses were then repeated to test for potential interactions between sex and each of the CBCL problem scales. There were no significant sex-by-behavior interactions across imaging modalities (**Table S2**).

#### Associations with Demographic Variables: Age, Sex and Cognitive Performance

Increased cognitive performance was consistently linked with increased odds of passing motion quality control across several imaging series including task-based fMRI: there was a 2% increase (*p_FDR_ =* 6.92e-19) for the SST task, 1% increase (*p_FDR_ =* 0.0009) for the MID task, and 3% increase (*p_FDR_ =* 1.71e-64) for the EN-Back task (**Table 2**). Male youth were less likely to pass the motion quality control than female youth for resting-state and the MID task with decreases in likelihood of 36.1% (*p_FDR_ =* 4.94e-11) and 22.1% (*p_FDR_ =* 3.94e-05), respectively (**Table 2**). Increased age was significantly associated with greater likelihood of passing motion quality control for resting-state, SST, and EN-Back tasks with increases of 3% (*p_FDR_ =* 2.53e-09), 1% (*p_FDR_ =* 0.0004), and 3% (*p_FDR_* = 2.83e-13), respectively (**Table 2**).

### Transdiagnostic Domains of Behavior and In-Scanner Motion

Significant associations were observed between attention-related behavioral problems and in-scanner motion for all imaging modalities (**Table 3**; **Figure 3**). That is, there was a positive association between greater severity of attention problems and greater motion during resting-state (*p_FDR_* = 3.96e-06), diffusion MRI (*p_FDR_* = 0.01), and all task fMRI series (all *p*s < 0.003) (**Table 3**). Of note, the greatest effects of attention problems on motion were observed for task-based fMRI, particularly for the EN-Back task (see **Table 3** for Cohen’s d effect size estimates). Regarding disruptive behavior and internalizing problems, significant associations were observed for resting-state and the EN-Back task (all *p*s < 0.01) (**Table 3**; **Figure 3**). Increased severity of disruptive behavior problems was associated with increased motion during resting-state (*p_FDR_* = 0.003) and the EN-Back task (*p_FDR_* = 0.003), while increased internalizing problem severity was associated with decreased motion during resting-state (*p* = 4.62e-05) and the EN-Back task (*p_FDR_* = 0.003). For the interested reader and to facilitate interpretation of findings, we also provide additional detail in the **Supplemental Results** regarding unit increases of motion associated with symptom domains.

**Figure 3.**
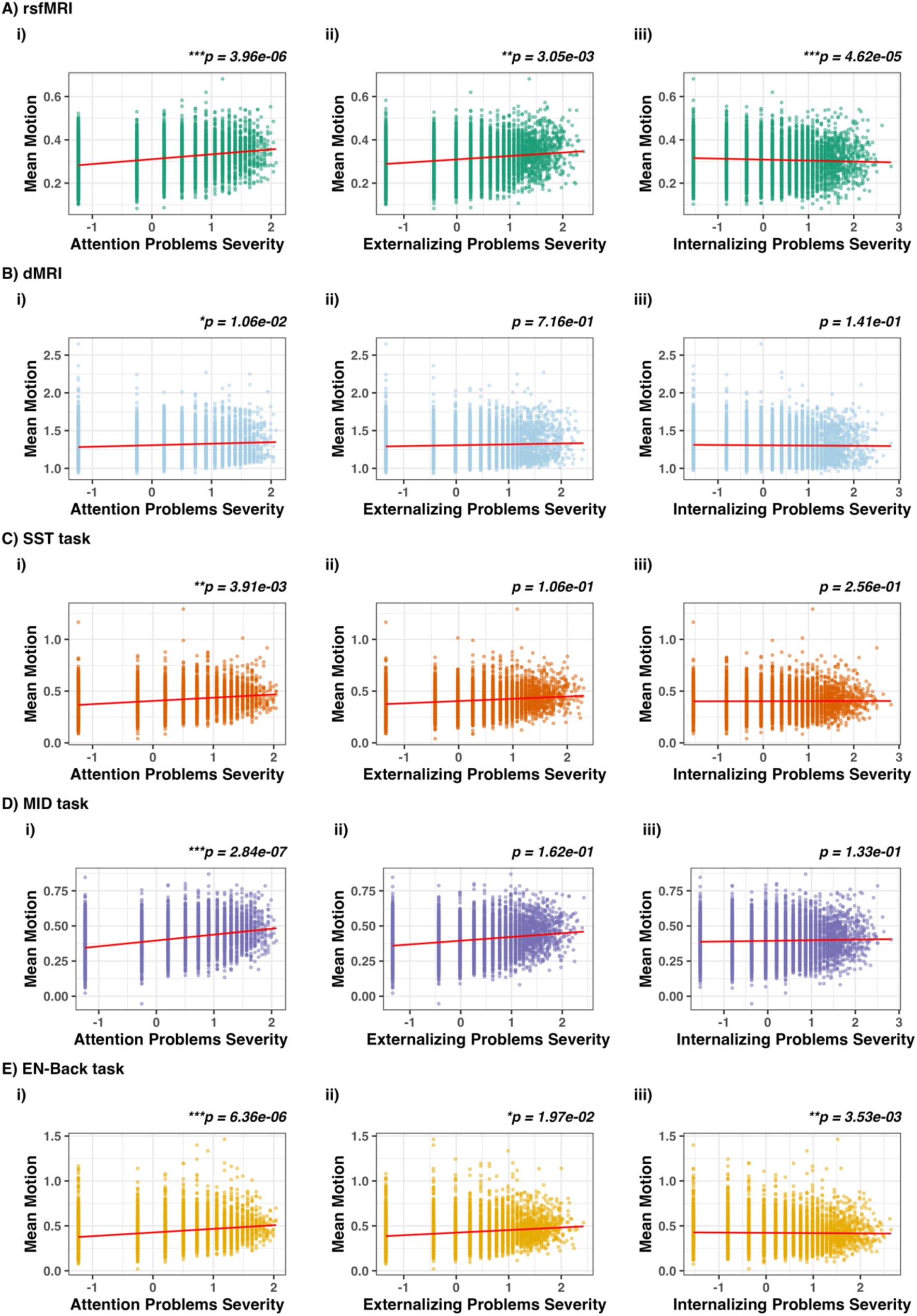
Severity of transdiagnostic symptom domains in youths are associated with motion during functional and structural imaging. Scatterplots depict results of linear mixed-effects models (see Table 3) for CBCL Attention, Externalizing, and Internalizing Problems scores for resting-state fMRI (A), diffusion MRI (B), Stop Signal Task (SST) (C), Monetary Incentive Delay (MID) task (D), and the emotional version of the n-back task (EN-Back) (E). The red trendline line represents the regression line based on the linear mixed-effects models fit. Statistical significance is denoted by asterisks: ***p < 0.001, **p < 0.01, and *p < 0.05.

**Table 3.** Results of Linear Mixed-Effects Models Predicting Motion and Domains of Transdiagnostic Symptoms.

|  | Estimate | Std.<br>Error | Cohen's D | t value | p | p<sub>FDR</sub> | 95% CI |  |
| --- | --- | --- | --- | --- | --- | --- | --- | --- |
|  |  |  |  |  |  |  | 2.5% | 2.5% |
| Resting-state fMRI |  |  |  |  |  |  |  |  |
| Intercept | 0.823 | 0.046 | 0.53 | 17.9 | 6.35e-69 | 3.49e-67 | 0.73 | 0.91 |
| Age | -0.003 | 0.0004 | -0.22 | -9.44 | 5.05e-21 | 3.97e-20 | -0.004 | -0.003 |
| Male | 0.028 | 0.005 | 0.12 | 5.14 | 2.81e-07 | 7.36e-07 | 0.02 | 0.04 |
| Black | 0.052 | 0.009 | 0.14 | 5.60 | 2.2e-08 | 6.71e-08 | 0.03 | 0.07 |
| Asian | 0.029 | 0.020 | 0.03 | 1.47 | 0.142 | 0.162 | -0.01 | 0.07 |
| Other | 0.021 | 0.009 | 0.06 | 2.31 | 0.021 | 0.0283 | 0.003 | 0.04 |
| Hispanic | 0.051 | 0.008 | 0.16 | 6.23 | 4.93e-10 | 1.69e-09 | 0.04 | 0.07 |
| Cognition | -0.001 | 0.0002 | -0.18 | -7.82 | 6.2e-15 | 2.84e-14 | -0.002 | -0.001 |
| CBCL Attention | 0.017 | 0.004 | 0.11 | 4.78 | 1.8e-06 | **3.96e-06** | 0.01 | 0.02 |
| CBCL Externalizing | 0.011 | 0.004 | 0.07 | 3.14 | 0.002 | **0.00305** | 0.004 | 0.02 |
| CBCL Internalizing | -0.014 | 0.003 | -0.09 | -4.24 | 2.27e-05 | **4.62e-05** | -0.02 | -0.01 |
| Diffusion MRI |  |  |  |  |  |  |  |  |
| Intercept | 1.73 | 0.097 | 1.65 | 17.7 | 4.22e-54 | 7.73e-53 | 1.54 | 1.92 |
| Age | -0.002 | 0.0007 | -0.066 | -2.81 | 0.00501 | 0.00744 | -0.003 | -0.0006 |
| Male | 0.008 | 0.011 | 0.018 | 0.77 | 0.439 | 0.465 | -0.01 | 0.03 |
| Black | 0.061 | 0.018 | 0.087 | 3.41 | 0.00065 | 0.00119 | 0.03 | 0.09 |
| Asian | 0.062 | 0.040 | 0.038 | 1.56 | 0.118 | 0.141 | -0.02 | 0.14 |
| Other | 0.030 | 0.018 | 0.043 | 1.64 | 0.1 | 0.125 | -0.01 | 0.07 |
| Hispanic | 0.046 | 0.016 | 0.073 | 2.87 | 0.00406 | 0.0062 | 0.02 | 0.08 |
| Cognition | -0.002 | 0.0003 | -0.128 | -5.38 | 7.56e-08 | 2.19e-07 | -0.002 | -0.001 |
| CBCL Attention | 0.018 | 0.007 | 0.063 | 2.68 | 0.0073 | **0.0106** | 0.005 | 0.03 |
| CBCL Externalizing | 0.003 | 0.007 | 0.009 | 0.38 | 0.703 | 0.716 | -0.01 | 0.02 |
| CBCL Internalizing | -0.010 | 0.007 | -0.036 | -1.55 | 0.12 | 0.141 | -0.02 | 0.003 |
| SST task |  |  |  |  |  |  |  |  |
| Intercept | 1.09 | 0.079 | 0.41 | 13.8 | 2.73e-42 | 3e-41 | 0.94 | 1.25 |
| Age | -0.004 | 0.0006 | -0.154 | -6.49 | 9.02e-11 | 3.31e-10 | -0.005 | -0.003 |
| Male | 0.054 | 0.009 | 0.138 | 5.72 | 1.08e-08 | 3.5e-08 | 0.04 | 0.07 |
| Black | 0.051 | 0.017 | 0.078 | 3.07 | 0.00215 | 0.00359 | 0.02 | 0.08 |
| Asian | 0.007 | 0.033 | 0.006 | 0.23 | 0.821 | 0.821 | -0.06 | 0.07 |
| Other | 0.039 | 0.016 | 0.062 | 2.37 | 0.0179 | 0.0246 | 0.007 | 0.07 |
| Hispanic | 0.055 | 0.014 | 0.108 | 3.85 | 0.000117 | 0.000222 | 0.03 | 0.08 |
| Cognition | -0.002 | 0.0003 | -0.197 | -8.21 | 2.71e-16 | 1.35e-15 | -0.003 | -0.002 |
| CBCL Attention | 0.018 | 0.006 | 0.0721 | 3.03 | 0.00242 | **0.00391** | 0.006 | 0.03 |
| CBCL Externalizing | 0.011 | 0.006 | 0.0412 | 1.73 | 0.0828 | 0.106 | -0.001 | 0.02 |
| CBCL Internalizing | -0.007 | 0.006 | -0.028 | -1.18 | 0.238 | 0.256 | -0.02 | 0.005 |
| **MID task** |  |  |  |  |  |  |  |  |
| Intercept | 1.16 | 0.069 | 0.486 | 16.7 | 3.92e-61 | 1.08e-59 | 1.02 | 1.3 |
| Age | -0.005 | 0.0005 | -0.199 | -8.69 | 4.54e-18 | 2.77e-17 | -0.006 | -0.004 |
| Male | 0.070 | 0.008 | 0.198 | 8.52 | 1.95e-17 | 1.07e-16 | 0.05 | 0.09 |
| Black | 0.069 | 0.014 | 0.121 | 4.88 | 1.07e-06 | 2.45e-06 | 0.04 | 0.09 |
| Asian | -0.021 | 0.030 | -0.017 | -0.71 | 0.479 | 0.497 | -0.080 | 0.04 |
| Other | 0.025 | 0.014 | 0.045 | 1.76 | 0.0792 | 0.104 | -0.003 | 0.05 |
| Hispanic | 0.061 | 0.012 | 0.135 | 4.93 | 8.27e-07 | 1.98e-06 | 0.04 | 0.09 |
| Cognition | -0.003 | 0.0003 | -0.229 | -9.85 | 9.86e-23 | 9.04e-22 | -0.003 | -0.002 |
| CBCL Attention | 0.028 | 0.005 | 0.122 | 5.33 | 1.03e-07 | **2.84e-07** | 0.02 | 0.04 |
| CBCL Externalizing | 0.008 | 0.006 | 0.034 | 1.46 | 0.144 | 0.162 | -0.003 | 0.02 |
| CBCL Internalizing | -0.008 | 0.005 | -0.037 | -1.6 | 0.109 | 0.133 | -0.02 | 0.002 |
| **EN-Back task** |  |  |  |  |  |  |  |  |
| Intercept | 1.19 | 0.081 | 0.431 | 14.6 | 1.79e-47 | 2.46e-46 | 1.03 | 1.35 |
| Age | -0.005 | 0.0006 | -0.176 | -7.2 | 6.52e-13 | 2.56e-12 | -0.006 | -0.003 |
| Male | 0.074 | 0.010 | 0.187 | 7.69 | 1.67e-14 | 7.08e-14 | 0.06 | 0.09 |
| Black | 0.072 | 0.017 | 0.108 | 4.17 | 3.06e-05 | 6.02e-05 | 0.04 | 0.11 |
| Asian | 0.047 | 0.034 | 0.035 | 1.39 | 0.164 | 0.18 | -0.02 | 0.113 |
| Other | 0.050 | 0.017 | 0.081 | 3.02 | 0.00257 | 0.00403 | 0.02 | 0.08 |
| Hispanic | 0.073 | 0.015 | 0.146 | 5.01 | 5.78e-07 | 1.44e-06 | 0.05 | 0.10 |
| Cognition | -0.003 | 0.0003 | -0.216 | -8.87 | 9.01e-19 | 6.19e-18 | -0.003 | -0.002 |
| CBCL Attention | 0.029 | 0.006 | 0.114 | 4.67 | 3e-06 | **6.36e-06** | 0.02 | 0.04 |
| CBCL Externalizing | 0.016 | 0.006 | 0.059 | 2.46 | 0.014 | **0.0197** | 0.003 | 0.03 |
| CBCL Internalizing | -0.018 | 0.006 | -0.075 | -3.08 | 0.00205 | **0.00353** | -0.03 | -0.007 |

Note: CBCL, Child Behavior Checklist; EN-Back, Emotional N-Back Task; MID, Monetary Incentive Delay Task; SST, Stop Signal Task. General Cognition is measured by the NIH Toolbox age corrected scores (35).

#### Interactions with Sex and Symptom Domains

Analyses were repeated to test for potential interactions between sex and each of the CBCL problem scales. There were no significant sex-by-behavior interactions across imaging modalities (**Table S3**).

#### Associations with Demographic Variables: Age, Sex and Cognitive Performance

Increased cognitive performance was consistently associated with decreased motion across all imaging modalities (all *p*s<0.0001) (**Table 3**). For male youth, there were also significant associations with increased motion during resting-state (*p_FDR_* =7.36e-07), SST (*p_FDR_* = 3.5e-08), MID (*p_FDR_* = 1.07e-16), and EN-Back (*p_FDR_* = 7.08e-14) tasks relative to female youth (**Table 3**). Increased age was significantly associated with decreased motion for all modalities (all *p*s <0.007) (**Table 3**).

### Follow-up Supplemental Analyses of Quality Control and In-Scanner Motion Across Imaging Modalities

As a supplemental analysis, we examined differences in the rates of passing quality control as well as mean motion between each modality. There were significant differences in quality control pass rates across the imaging modalities (*p* < 2e-16): rsfMRI = MID > diffusion MRI > SST > EN-Back (**Figure 4** and **Supplemental Results**). There were also significant differences in mean FD across the imaging modalities (*p* < 2e-16): diffusion MRI > SST > EN-Back > MID > resting-state (**Figure 5** and **Supplemental Results**).

**Figure 4.**
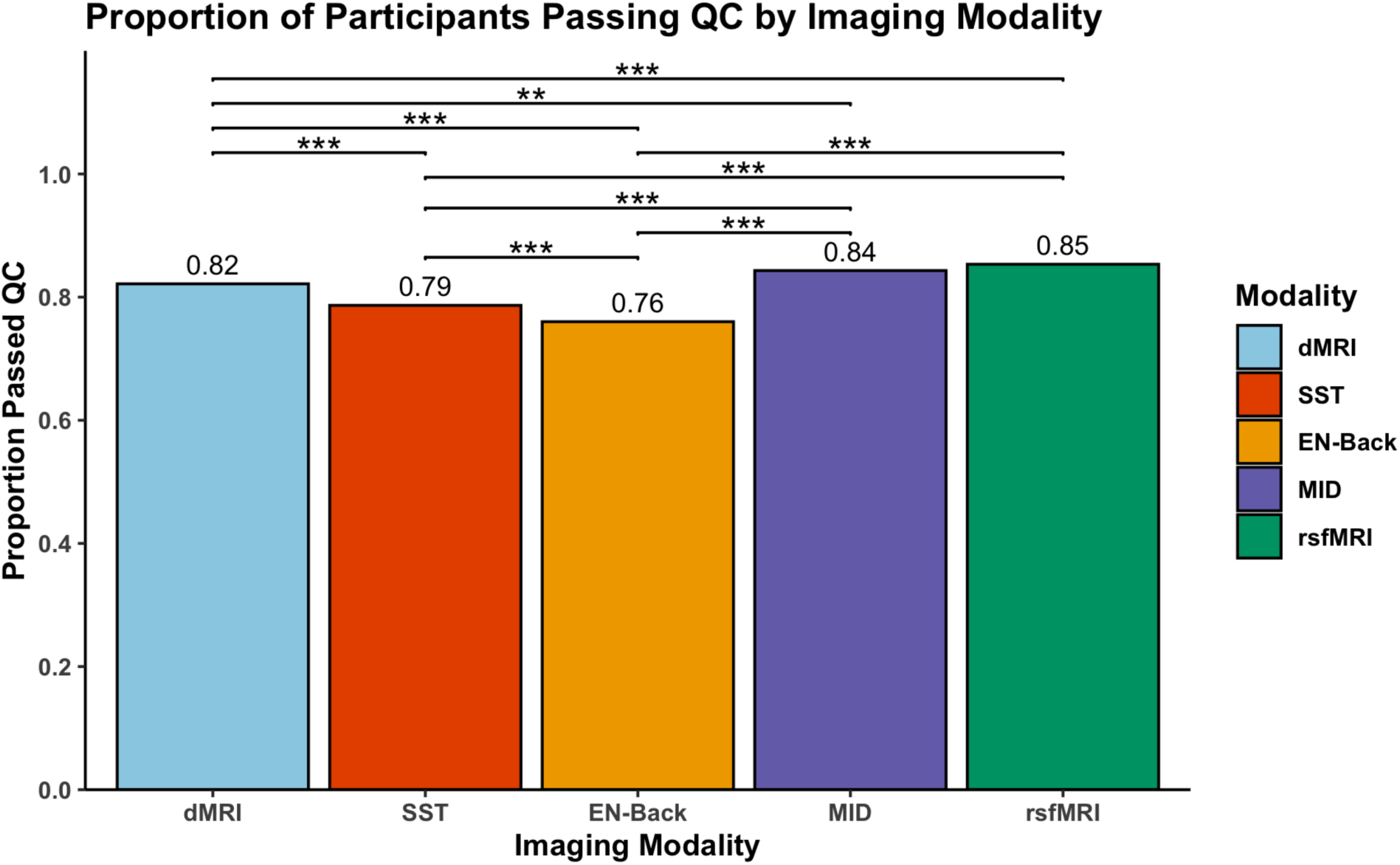
Proportion of participants passing quality control by imaging modality. The bar plot represents the proportion of participants who passed quality control (QC) across five different imaging modalities: resting-state fMRI (rsfMRI), diffusion MRI (dMRI), Stop Signal Task (SST), Monetary Incentive Delay (MID) task, and the emotional version of the n-back task (EN-Back). Statistical significance is denoted by asterisks: ***p < 0.001, **p < 0.01, *p < 0.05.

**Figure 5.**
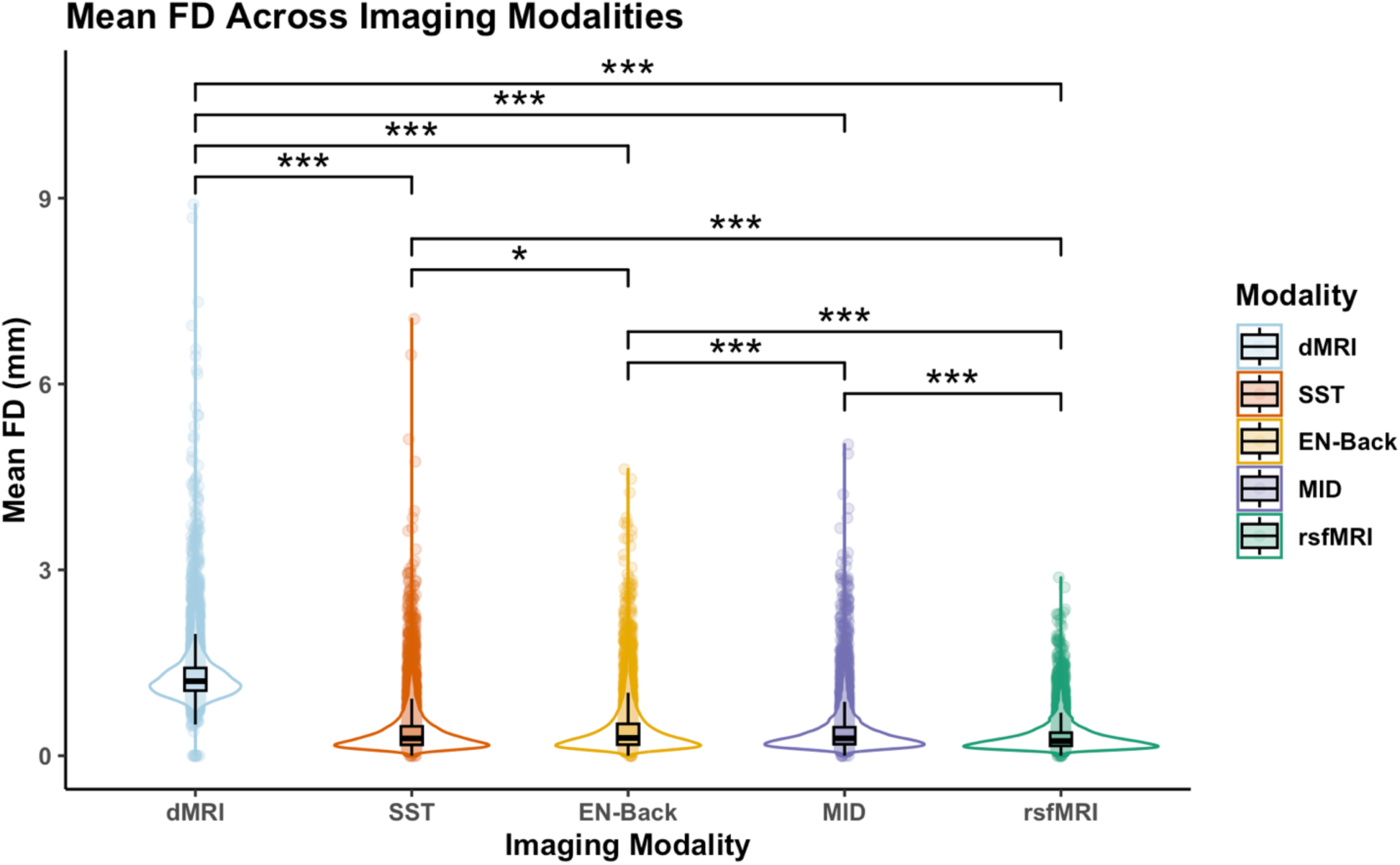
Motion in youths varies across functional and structural imaging modalities. Violin plots represent the mean framewise displacement (FD) for each imaging modality: resting-state fMRI (rsfMRI), diffusion MRI (dMRI), Stop Signal Task (SST), Monetary Incentive Delay (MID) task, and the emotional version of the n-back task (EN-Back). Statistical significance is denoted by asterisks: ***p < 0.001, **p < 0.01, *p < 0.05.

## Discussion

The current study examined the relationships between transdiagnostic symptom domains and head motion during multimodal imaging in children. Three main findings emerged. First, greater attention and disruptive behavior problems in youths are linked to reduced likelihood of passing motion quality control, while greater internalizing symptoms are linked to greater likelihood of passing motion quality control across imaging modalities. Second, greater severity of transdiagnostic symptom domains of attention and disruptive behavior problems was associated with greater in-scanner motion across modalities. However, greater internalizing symptom severity was associated with reduced motion. Third, supplemental analyses indicated no interaction of sex differences in scan motion across transdiagnostic symptom domains, suggesting similar effects of motion across girls and boys related to attention, internalizing and disruptive behavior problem severities. Consistent with prior work, findings also indicated associations between greater motion with younger age and reduced cognitive performance, as well as greater motion in boys vs girls. This study suggests that severity of symptom domains may be linked to motion in youths, which has implications for developing robust and reliable methods for mitigating motion in translational developmental neuroscience as well as considerations for enhancing accessibility of imaging protocols for youths with varying clinical phenotypes.

### Transdiagnostic Domains of Behavior and Imaging Quality Control

Significant associations were found between transdiagnostic domains of behavior and passing motion quality control. Particularly, children with increased attention-related difficulties exhibited significantly lower likelihood of passing motion quality control across all imaging modalities. Additionally, increased severity of disruptive behavior problems was also associated with a decrease (9.4%) in the likelihood of passing motion quality control during the EN-back task. These findings are consistent with previous results indicating that increased attention-related problems may impact scanning for youths with a greater likelihood of motion and artifact, thereby impacting data retention (7). Additionally, the most substantial decrease in passing rates was observed in T1-weighted scans (25.8%) followed by resting-state (20.2%), diffusion MRI (19.9%), SST task (15.4%), MID task (10.1%), and the EN-Back task (10.8%). The variation in motion across different imaging modalities could suggest that tasks requiring minimal engagement (e.g., T1-weighted scans, resting-state, and/or diffusion MRI), may be more susceptible to motion in children with elevated severity of attention-related challenges. In contrast, the potentially more engaging nature of task-based fMRI (e.g., SST, MID, and EN-Back tasks), likely facilitate engagement during scanning for children with attention-related challenges, which aligns with the observed higher pass rates of quality control in the current study. This is also consistent with the notion that head motion tends to be lower when youths are actively engaged in cognitive and/or language tasks (2, 27). However, it is important to note that studies have also reported greater levels of scanner motion related to more demanding cognitive control tasks, which may increase the challenge in monitoring motion during stimuli, distractions, and/or sustained attention (1, 31, 48).

Our finding of increased motion linked to increased severity of attention and disruptive behavior problems aligns with prior work suggesting that youths with elevated severity of these symptom domains are more prone to scanner movement relative to age-matched controls (2, 49, 50). Associations have also been reported between high-motion subgroups and total CBCL Problem Behaviors relative to low-motion subgroups (48) as well as between motion in youths with neurodevelopmental conditions (including ASD and ADHD) vs neurotypical groups (6). The EN-Back task, which involves working memory and requires sustained attention and behavioral control, may pose greater challenges for children with behavioral difficulties related to attention and disruptive behavior problems, thereby affecting the ability to remain still during scanning. Interestingly, increased internalizing problem severity was linked with an increase (10%) in the likelihood of passing motion quality control checks during the SST task. This finding suggests that children with greater severity of internalizing symptoms, such as anxiety, may exhibit reduced movement during task-based fMRI. The nature of the SST task, which requires participants to inhibit ongoing actions, provides a measure of the speed of these inhibitory processes. Here, increased internalizing symptoms could reflect anxiety-related symptoms such as fearing negative evaluation, which may impact fMRI task compliance (51). Alternately, greater motion associated with attention and disruptive behavior problems during task-based fMRI could also reflect difficulties in cognitive control processes (e.g., inhibitory control, cognitive flexibility), which may not necessarily emerge as areas of cognitive difficulty for youths with internalizing problems (52).

### Transdiagnostic Domains of Behavior and In-Scanner Motion

Associations were also observed between transdiagnostic symptom domains and in-scanner head motion (indexed using mean FD) across functional and structural imaging. Children with increased attention-related behavioral challenges showed increased motion across all modalities, particularly during task-based fMRI. These findings are consistent with prior work suggesting an association between attention-related symptoms and increased motion in youths, even despite motion scrubbing or censoring approaches (7, 24). The few existing studies exploring scanner motion in clinical groups vs unaffected controls have also reported increased motion in individuals with ASD, ADHD, schizophrenia, and bipolar disorder vs unaffected controls (2, 4). These findings could suggest that children with greater severity of attention-related difficulties are more prone to movement during fMRI tasks requiring active engagement that tap into cognitive control processes. In support of this idea, the use of engaging movies during scanning in younger children has been shown to reduce motion effects, particularly compared to resting-state acquired with eyes open and attention cued to a fixation cross (1, 48, 53). However, it is also possible that within- and between-network connections in sensorimotor, visual and cognitive control circuits may be more affected and altered by movie watching relative to conventional resting-state paradigms involving a fixation cross-hair (1). Our findings also suggest that during task-based fMRI, children tend to show relatively greater tolerance to motion (i.e., lower motion). It is possible that tasks such as SST or variations thereof may provide an optimal balance for younger children that facilitates cognitive engagement during a moderately- or fast-paced paradigm with a requirement for frequent responses and attending (24, 32). While head motion tends to be lower when participants are actively engaged in a cognitive task, certain task conditions may show comparable motion depending on the cognitive and/or language demands required (2, 27, 54). Additionally, fMRI tasks that integrate a combination of visual, auditory, and manual response components may help to reduce motion in youths during scanning (27). Alternatively, our findings may also suggest that motion could be associated with differences in subject groups (e.g., clinical groups vs unaffected controls) rather than cognitive processes associated with task properties (22, 55).

Our results also indicate that increased severity of disruptive behavior problems is associated with increased head motion during resting-state fMRI and the EN-Back task, but the opposite pattern was observed for internalizing problems: that is, increased severity of internalizing symptoms was associated with decreased head motion. These findings are consistent with previous studies indicating associations between motion and behavioral or symptom severity in youths (4, 48), particularly when motion is modeled as a continuous variable. Similarly, Nebel et al. (4) showed that autistic children were more likely to be excluded than unaffected, neurotypical children regardless of lenient vs conversative motion criterion; notably, the subsample of autistic children with data passing quality control criterion tended to be older, have milder social impairments, better motor control, and higher cognitive performance than the total sample. However, few studies have unpacked transdiagnostic domains of behavior and/or prior studies have been limited by small, community-based samples. Here, we examined correlates of motion across functional and structural imaging modalities with transdiagnostic domains of behavior related to disruptive behavior and internalizing symptoms (modeled as continuous variables) to differentiate unique patterns of motion among symptom domains. It is also possible that differences in motion may compound direct comparisons of measures of functional connectivity among clinical groups vs unaffected controls, considering potential between-group differences in motion (2). For instance, in imaging samples combining clinical and community participants, exclusion of high-motion subgroups may impact biases toward the selection of children with lower overall symptom severities (e.g., social functioning, attention, and disruptive behavior problems), which may differ from the initial target population and reduce statistical power for identifying between-group differences in clinical vs unaffected control groups: that is, the severity of clinical phenotypes in low-motion youths may be more similar to unaffected controls (4).

### Limitations

There are limitations to acknowledge. First, we did not investigate the influence of movement or data scrubbing on image quality, brain activation or functional connectivity. While these aspects were beyond the scope of the current study, future work will be important to examine associations between motion, transdiagnostic symptom domains, and measures of functional connectivity. Second, ABCD Study participants primarily consisted of a community-based sample of children with varying levels of externalizing and internalizing symptoms. This may limit the generalizability of our findings to clinical groups or youths with more severe symptomatology. Thus, future work testing replication of findings in clinical groups will be important. Nonetheless, the ABCD Study sample provides heterogeneity and diversity both demographically and across a range of clinical phenotypes. Third, the variability in fMRI tasks, behavioral measures, and acquisition protocols across different research groups beyond ABCD Study sites may limit the generalizability of our findings to other research and/or clinical settings. However, it is important to emphasize that the ABCD Study protocol was designed to optimize harmonization of sequence acquisition protocols across the 21 sites and uses well-established measures of child psychopathology (CBCL) (32, 33). Fourth, the age range of participants was narrow (9-10 years in the first wave), and future longitudinal work will be needed to understand the stability of motion across development, particularly given the potential heritable contributions of motion stability throughout development (24). However, we opted to use a narrow age range to limit developmental heterogeneity of the sample as a potential confound in analyses.

### Recommendations for Translational Developmental Neuroimaging

Increasing accessibility and feasibility of scanning protocols for children and adolescents will be important for enhancing the heterogeneity of community as well as clinical samples and for increasing generalizability of imaging findings. Here, we also provide recommendations and considerations that could potentially help mitigate motion and accommodate inter-individual differences in behavioral, cognitive, and language abilities in youth to facilitate accessibility of scanning in pediatric populations. First, the association between transdiagnostic domains of symptoms and motion during scanning underscores the necessity for customized imaging protocols that can accommodate children with various behavioral challenges. For example, children with elevated attention-related difficulties, anxiety and/or sensory sensitivities might benefit from shorter scanning sessions or integrating engaging, interactive tasks that help maintain focus and reduce motion. When developing fMRI paradigms, another consideration is the inclusion of audio and visual cues together with button presses or explicit responses to the stimuli (vs a passive viewing task), which may help to reduce motion in children (1, 27, 48). Second, our findings also emphasize the need for continued advancement and integration of motion correction methods. Elevated motion is also shown to decrease reliability and stability estimates within cognitive control circuitry during ABCD task-based fMRI paradigms when comparing subgroups of children in the highest vs lowest movement quartiles (3). Thus, developing robust computational models and algorithms that address motion without compromising data and timeseries integrity will continue to be important to improve imaging data quality and retention in developmental populations. For instance, the use of real-time motion detection, or framewise integrated real-time MRI monitoring (FIRMM) software, in which participants receive feedback during scanning, is a highly valuable advancement in mitigating motion effects and increasing data retention that was implemented in 15 out of 21 ABCD Study sites (1, 6, 33). However, even with the use of FIRMM, Marek et al. (56) excluded 40% and Marek et al. (57) excluded 60% of participants with resting-state fMRI data to ensure low-motion samples. Recent work has also explored machine learning models leveraging a Doubly Robust Targeted Minimum Loss based Estimation (DRTMLE) approach, which treats excluded resting-state scans as a missing data problem to address bias concerns for functional connectivity (i.e., exclusion of high-motion participants may introduce bias by systematically altering the study population) (4). In addition to prospectively correcting for motion during data acquisition and preprocessing, we also recommend regressing motion estimates from between-subject analyses (e.g., group-level contrasts of clinical groups vs unaffected controls). Third, implementing a mock scan training session, prior to the actual scan, may be helpful in acclimating children to the scanner environment and reducing motion. Mock scanners equipped with real-time feedback, such as the MoTrak Head Motion Tracking System (Psychology Software Tools, 2017), provide the participant with visual cues for training (i.e., cursor within a target region) when motion exceeds a threshold (e.g., >3 mm translational motion will pause a movie or video). In our research group, the use of mock scans with real-time feedback has received positive feedback from participants and has shown promising results with similar motion during scanning between clinical and unaffected control groups (58, 59). Other considerations include extended and/or multiple mock scanner sessions based on the needs of the participant. Fourth, providing an option of breaking up imaging acquisition into multiple sessions on the same day as needed, as implemented in the ABCD Study (32), and/or options for movement breaks during a scanning session (depending on the child’s needs) may help mitigate motion effects in youths (54). For instance, given that motion has been shown to increase with ongoing session length (54, 60), participants could be offered minor movement breaks within the scanner between transitions of each scan acquisition and/or the option to divide longer data acquisitions into two shorter sessions on the same day, similar to the ABCD Study (32). Lastly, teaching simple and feasible emotion regulation strategies that might not interfere with fMRI paradigms and targeted cognitive processes could be helpful in reducing scan-related anxiety. For instance, reviewing simple behavioral and cognitive strategies prior to the scan including mental imagery and positive self-statements (e.g., “I can do this”, “do your best”) may equip participants with fundamental regulation abilities to build confidence while optimizing motion reduction during scanning. Cognitive strategies may also be paired with behavioral training or rehearsal during the mock scan to reinforce skill acquisition and generalization to the actual scanner. Additionally, given that motion has been shown to be relatively stable within participants (vs between participants) with a potential genetic influence (22, 24, 25, 60, 61), individualizing approaches for motion reduction and scanner training will be essential.

## Conclusion

Findings from the current study suggest that transdiagnostic symptom domains of attention and disruptive behavior problems are associated with increased motion in youths. In contrast, elevated internalizing problems are associated with reduced motion. Future work is needed to continue to develop robust computational approaches and behavioral methods to mitigate motion effects in youths. Enhancing accessibility of neuroimaging protocols in pediatric populations with consideration for accommodating a range of symptom severities and behavioral challenges in children will have implications for advancing development of robust, reliable and generalizable brain biomarkers for child mental health.

## Acknowledgments

K.I. is supported by the National Institute of Mental Health (K23-MH128451). This work was supported by National Center for Advancing Translational Sciences grant KL2 TR001862 (K.I.) and TL1 TR001864 (K.I.), a Yale Child Study Center Junior Faculty Development Pilot Award (K.I.), and the Yale Child Study Center Translational Developmental Neuroscience Training Program (T32 MH18268) (K.I.). K.H. and Z.L. are supported by the Horstmann Scholarship from the Yale University School of Public Health.

## ABCD acknowledgement

A portion of the data used in the preparation of this article were obtained from the Adolescent Brain Cognitive Development (ABCD) Study (https://abcdstudy.org), held in the NIMH Data Archive (NDA). This is a multisite, longitudinal study designed to recruit more than 10,000 children ages 9-10 and follow them over 10 years into early adulthood. The ABCD Study® is supported by the National Institutes of Health and additional federal partners under award numbers U01DA041022, U01DA041028, U01DA041048, U01DA041089, U01DA041106, U01DA041117, U01DA041120, U01DA041134, U01DA041148, U01DA041156, U01DA041174, U24DA041123, U24DA041147, U01DA041093, and U01DA041025. A full list of supporters is available at https://abcdstudy.org/federal-partners.html. A listing of participating sites and a complete listing of the study investigators can be found at https://abcdstudy.org/consortium_members/. ABCD consortium investigators designed and implemented the study and/or provided data but did not necessarily participate in analysis or writing of this report. This manuscript reflects the views of the authors and may not reflect the opinions or views of the NIH or ABCD consortium investigators. The ABCD data repository grows and changes over time. The ABCD data used in this report came from NDA 4.0 release (DOI: <u>10.15154/z563-zd24</u>). We are grateful to the study participants for their time and participation.

## Preprint Servers

A version of this manuscript was posted as a preprint on *bioRxiv*. The authors retain full copyright.

