## Supplemental Information for "Transdiagnostic Symptom Domains are Associated with Head Motion During Multimodal Imaging in Children"

Karim Ibrahim, Psy.D.

Assistant Professor

Yale University School of Medicine

Child Study Center

230 South Frontage Road

New Haven, CT 06520

**Supplemental Methods:**

**ABCD study inclusion variables/flags**

The inclusion criteria for each imaging modality are aligned with the “ABCD Release 4.0 release notes” available at DOI 10.15154/1523041: “NDA 4.0 MRI Quality Control Recommended Inclusion”. For more detail regarding the ABCD study structural (T1, diffusion MRI), resting-state, and task sequences and acquisition parameters, see Casey et al. (1) and Hagler et al. (2). We provide a summary from the release notes of these inclusion criteria as follows:

**T1-weighted (variable *imgincl_t1w_include* = 1):** Must pass both raw QC for T1 and FreeSurfer QC. Derived results must exist.

**Diffusion MRI (variable *imgincl_dmri_include* = 1):** Must pass raw QC and T1 QC, with acceptable repetitions. B0 unwarp must be available. FreeSurfer QC and manual post-processing QC must be successful. Registration to T1w must be less than 17. Dorsal cutoff score must be less than 47, and ventral cutoff score less than 54. Derived results must exist.

**Resting-state fMRI (variable *imgincl_rsfmri_include* = 1):** Must pass raw QC and T1 QC, with more than 375 frames post-censoring. B0 unwarp must be available. FreeSurfer QC and manual post-processing QC must be successful. Registration to T1w must be less than 19. Dorsal cutoff score must be less than 65, and ventral cutoff score less than 60. Derived results must exist.

**MID** **task (variable *imgincl_mid_include* = 1):** Must pass raw QC and T1 QC. Behavioral performance must be acceptable with more than 200 degrees of freedom. B0 unwarp must be available. FreeSurfer QC and manual post-processing QC must be successful. Registration to T1w must be less than 19. Dorsal cutoff score must be less than 65, and ventral cutoff score less than 60. Derived results must exist.

**EN-Back** **task** **(variable *imgincl_nback_include* = 1):** Must pass raw QC and T1 QC. Behavioral performance must be acceptable with more than 200 degrees of freedom. B0 unwarp must be available. FreeSurfer QC and manual post-processing QC must be successful. Registration to T1w must be less than 19. Dorsal cutoff score must be less than 65, and ventral cutoff score less than 60. Derived results must exist.

**SST (variable *imgincl_sst_include* = 1):** Must pass raw QC and T1 QC. Behavioral performance must be acceptable with no glitches and more than 200 degrees of freedom. B0 unwarp must be available. FreeSurfer QC and manual post-processing QC must be successful. Registration to T1w must be less than 19. Dorsal cutoff score must be less than 65, and ventral cutoff score less than 60. Derived results must exist.

**Supplemental Results:**

**Unit Increases of Motion Associated with Symptom Domains:** To facilitate interpretation of findings, we provide additional detail regarding unit increases of motion related to each of the symptom domains. A single unit increase in the CBCL Attention Problems score corresponded to a 0.017 mm increase in motion during resting-state fMRI. This finding was consistent across modalities, with single unit increases associated with 0.018 mm increases during both diffusion MRI and the SST task, a 0.028 mm increase during the MID task, and a 0.029 mm increase during the EN-Back task. An increase in the CBCL Externalizing Problems score was significantly associated with a 0.012 mm increase in motion during resting-state and a 0.016 mm increase during the EN-Back task. Conversely, an increase in the CBCL Internalizing Problems score corresponded to a 0.014 mm decrease in motion during resting-state and a 0.018 mm decrease during the EN-Back task.

**Associations with Demographic Variables**: Results of logistic regression models showed that other demographic variables also demonstrated significant associations with passing motion quality control (**Table 2**). Black participants were less likely to pass motion quality control than White participants for the SST and EN-Back tasks with decreases in likelihood of 21.6% (*p* = 0.01) and 29.4% (*p* = 0.0001), respectively. Results of linear mixed-effects models revealed that race showed statistically significant effects across all imaging modalities (**Table 3**). Black participants showed significant positive associations with increased motion across imaging modalities compared to White participants (all *P*s<0.001). There were also significant associations for ethnicity (i.e., Hispanic vs non-Hispanic) across all imaging series (all *P*s<0.006).

**Differences for Proportion of Participants Passing Quality Control Across Imaging Modalities.** As a supplemental analysis, we also evaluated the rates of passing quality control across imaging modalities. Results of a Pearson’s Chi-squared test demonstrated significant variability in pass rates across modalities (χ² = 363.32, df = 4, *p* < 2.2e-16). Follow-up pairwise comparisons of proportions, adjusted for multiple testing using the Bonferroni method, revealed significant differences between several pairs of modalities (**Figure 5**). Significant differences were found between dMRI and rsfMRI (*p* = 7.3e-08), dMRI and MID (*p* = 0.00113), dMRI and EN-Back (*p* < 2e-16), dMRI and SST (*p* = 3.6e-08), rsfMRI and EN-Back (*p* < 2e-16), rsfMRI and SST (*p* < 2e-16), MID and EN-Back (*p* < 2e-16), MID and SST (*p* < 2e-16), as well as EN-Back and SST (*p* = 0.00022). In contrast, no significant difference was found between rsfMRI and MID (*p* = 0.56576).

**Differences for In-Scanner Motion Across Imaging Modalities.** As a supplemental analysis, we examined differences in motion between each modality. There were significant differences in mean FD across the imaging modalities (*F*(4, 36,761) = 8,653, *p* < 2e-16). diffusion MRI>SST > EN-Back > MID > resting-state. Post-hoc tests revealed significantly greater motion during diffusion MRI compared to other modalities (all *P*s < .001) (**Figure 4**): diffusion MRI (*M* = 1.31, *SD* = 0.48) vs SST (*M*= 0.40, *SD* = 0.40), EN-Back (*M* = 0.42, *SD* = 0.40), MID (*M* = 0.39, *SD* = 0.37), and resting-state (*M* = 0.31, *SD* = 0.24). Additionally, there was significantly greater motion during SST vs EN-Back (*p* = .019) and resting-state (*p* < .001). There was also significantly greater motion during the EN-Back vs MID and resting-state (all *P*s < .001), and greater motion during the MID task vs resting-state (p < .001).

**Table S1:** ABCD Study Site Data for Each Imaging Modality

| Site |  | T1-Weighted | |  | Resting-state fMRI | |  | Diffusion MRI | |  | SST Task | |  | MID Task | |  | EN-Back Task | |
| --- | --- | --- | --- | --- | --- | --- | --- | --- | --- | --- | --- | --- | --- | --- | --- | --- | --- | --- |
|  | Total | Pass | Fail |  | Pass | Fail |  | Pass | Fail |  | Pass | Fail |  | Pass | Fail |  | Pass | Fail |
| site01 | 280 | 96.07 | 3.93 |  | 83.57 | 16.43 |  | 0 | 100 |  | 79.29 | 20.71 |  | 86.43 | 13.57 |  | 75.71 | 24.29 |
| site02 | 463 | 99.35 | 0.65 |  | 95.03 | 4.97 |  | 97.41 | 2.59 |  | 87.47 | 12.53 |  | 89.20 | 10.80 |  | 83.80 | 16.20 |
| site03 | 470 | 100 | 0 |  | 94.26 | 5.74 |  | 99.15 | 0.85 |  | 84.26 | 15.74 |  | 91.28 | 8.72 |  | 77.45 | 22.55 |
| site04 | 656 | 87.65 | 12.35 |  | 80.79 | 19.21 |  | 82.47 | 17.53 |  | 69.82 | 30.18 |  | 75.46 | 24.54 |  | 66.16 | 33.84 |
| site05 | 310 | 98.71 | 1.29 |  | 87.10 | 12.90 |  | 98.39 | 1.61 |  | 80 | 20 |  | 83.55 | 16.45 |  | 71.94 | 28.06 |
| site06 | 434 | 98.85 | 1.15 |  | 87.10 | 12.90 |  | 96.54 | 3.46 |  | 82.95 | 17.05 |  | 87.56 | 12.44 |  | 81.34 | 18.66 |
| site07 | 255 | 97.25 | 2.75 |  | 84.71 | 15.29 |  | 94.90 | 5.10 |  | 75.29 | 24.71 |  | 87.45 | 12.55 |  | 76.86 | 23.14 |
| site08 | 223 | 94.17 | 5.83 |  | 86.10 | 13.90 |  | 84.30 | 15.70 |  | 80.72 | 19.28 |  | 81.61 | 18.39 |  | 74.44 | 25.56 |
| site09 | 318 | 97.48 | 2.52 |  | 87.42 | 12.58 |  | 95.60 | 4.40 |  | 80.82 | 19.18 |  | 89.62 | 10.38 |  | 76.42 | 23.58 |
| site10 | 583 | 91.77 | 8.23 |  | 77.70 | 22.30 |  | 83.70 | 16.30 |  | 68.10 | 31.90 |  | 77.02 | 22.98 |  | 66.21 | 33.79 |
| site11 | 397 | 98.74 | 1.26 |  | 83.38 | 16.62 |  | 97.73 | 2.27 |  | 76.32 | 23.68 |  | 87.15 | 12.85 |  | 73.30 | 26.70 |
| site12 | 520 | 97.50 | 2.50 |  | 83.27 | 16.73 |  | 95.00 | 5.00 |  | 74.04 | 25.96 |  | 85.00 | 15.00 |  | 75.38 | 24.62 |
| site13 | 526 | 85.36 | 14.64 |  | 77.95 | 22.05 |  | 79.28 | 20.72 |  | 67.11 | 32.89 |  | 75.86 | 24.14 |  | 65.97 | 34.03 |
| site14 | 459 | 98.91 | 1.09 |  | 86.49 | 13.51 |  | 97.17 | 2.83 |  | 82.57 | 17.43 |  | 89.32 | 10.68 |  | 82.57 | 17.43 |
| site15 | 260 | 95.38 | 4.62 |  | 73.46 | 26.54 |  | 90.38 | 9.62 |  | 67.31 | 32.69 |  | 73.46 | 26.54 |  | 61.92 | 38.08 |
| site16 | 916 | 99.34 | 0.66 |  | 96.51 | 3.49 |  | 98.03 | 1.97 |  | 91.16 | 8.84 |  | 91.92 | 8.08 |  | 90.83 | 9.17 |
| site17 | 408 | 96.57 | 3.43 |  | 75.74 | 24.26 |  | 0.00 | 100 |  | 84.31 | 15.69 |  | 85.54 | 14.46 |  | 77.45 | 22.55 |
| site18 | 301 | 91.03 | 8.97 |  | 80.07 | 19.93 |  | 86.38 | 13.62 |  | 70.76 | 29.24 |  | 73.42 | 26.58 |  | 73.75 | 26.25 |
| site19 | 319 | 97.81 | 2.19 |  | 82.45 | 17.55 |  | 0 | 100 |  | 82.45 | 17.55 |  | 85.27 | 14.73 |  | 73.98 | 26.02 |
| site20 | 515 | 98.25 | 1.75 |  | 86.99 | 13.01 |  | 93.59 | 6.41 |  | 80.00 | 20 |  | 85.44 | 14.56 |  | 78.25 | 21.75 |
| site21 | 408 | 97.30 | 2.70 |  | 87.50 | 12.50 |  | 94.36 | 5.64 |  | 77.70 | 22.30 |  | 82.84 | 17.16 |  | 76.23 | 23.77 |
| site22 | 24 | 91.67 | 8.33 |  | 87.50 | 12.50 |  | 91.67 | 8.33 |  | 87.50 | 12.50 |  | 79.17 | 20.83 |  | 83.33 | 16.67 |

**Table S2:** Results of Logistic Regression Models Predicting Scan Quality Control and Domains of Transdiagnostic Symptoms with Sex Interactions

| **Variables** | **Estimate** | **Std. Error** | ***z* value** | ***p*** | ***p*_FDR_** | **OR** | **95% CI** | |
| --- | --- | --- | --- | --- | --- | --- | --- | --- |
|  |  |  |  |  |  |  | **2.5 %** | **97.5 %** |
| **T1-Weighted** |  |  |  |  |  |  |  |  |
| Intercept | 1.7 | 0.942 | 1.81 | 0.0709 | 0.213 | 5.49 | 0.87 | 34.8 |
| Age | 0.012 | 0.007 | 1.67 | 0.094 | 0.255 | 1.01 | 0.99 | 1.03 |
| Male | -0.067 | 0.115 | -0.58 | 0.562 | 0.749 | 0.935 | 0.75 | 1.17 |
| Black | -0.221 | 0.171 | -1.29 | 0.197 | 0.425 | 0.802 | 0.57 | 1.12 |
| Asian | 0.072 | 0.163 | 0.44 | 0.658 | 0.789 | 1.07 | 0.78 | 1.48 |
| Other | -0.396 | 0.331 | -1.19 | 0.232 | 0.482 | 0.673 | 0.35 | 1.29 |
| Hispanic | -0.094 | 0.17 | -0.55 | 0.58 | 0.756 | 0.91 | 0.65 | 1.27 |
| Cognition | 0.007 | 0.003 | 1.98 | 0.048 | 0.161 | 1.01 | 1 | 1.01 |
| Attention | -0.133 | 0.103 | -1.29 | 0.197 | 0.425 | 0.875 | 0.72 | 1.07 |
| Externalizing | 0.16 | 0.108 | 1.48 | 0.138 | 0.322 | 1.17 | 0.95 | 1.45 |
| Internalizing | 0.067 | 0.099 | 0.68 | 0.498 | 0.71 | 1.07 | 0.88 | 1.3 |
| Sex-by-CBCL Attention | -0.32 | 0.145 | -2.21 | 0.0273 | 0.114 | 0.726 | 0.55 | 0.97 |
| Sex-by-CBCL Externalizing | -0.036 | 0.144 | -0.25 | 0.8 | 0.861 | 0.965 | 0.73 | 1.28 |
| Sex-by-CBCL Internalizing | 0.095 | 0.133 | 0.72 | 0.473 | 0.698 | 1.1 | 0.85 | 1.43 |
| **Resting-state fMRI** |  |  |  |  |  |  |  |  |
| Intercept | -2.25 | 0.547 | -4.12 | 3.84e-05 | 0.000358 | 0.105 | 0.04 | 0.31 |
| Age | 0.027 | 0.004 | 6.39 | 1.67e-10 | 2.33e-09 | 1.03 | 1.02 | 1.04 |
| Male | -0.44 | 0.065 | -6.75 | 1.46e-11 | 2.45e-10 | 0.644 | 0.57 | 0.73 |
| Black | -0.07 | 0.101 | -0.69 | 0.49 | 0.709 | 0.932 | 0.76 | 1.14 |
| Asian | -0.076 | 0.096 | -0.79 | 0.425 | 0.688 | 0.927 | 0.77 | 1.12 |
| Other | -0.432 | 0.206 | -2.1 | 0.0361 | 0.132 | 0.649 | 0.43 | 0.97 |
| Hispanic | -0.08 | 0.105 | -0.76 | 0.445 | 0.688 | 0.923 | 0.75 | 1.13 |
| Cognition | 0.012 | 0.002 | 5.98 | 2.2e-09 | 2.63e-08 | 1.01 | 1.01 | 1.02 |
| Attention | -0.133 | 0.0625 | -2.13 | 0.0333 | 0.127 | 0.875 | 0.77 | 0.99 |
| Externalizing | 0.021 | 0.066 | 0.32 | 0.751 | 0.849 | 1.02 | 0.89 | 1.16 |
| Internalizing | 0.045 | 0.06 | 0.76 | 0.45 | 0.688 | 1.05 | 0.93 | 1.18 |
| Sex-by-CBCL Attention | -0.161 | 0.083 | -1.94 | 0.053 | 0.165 | 0.851 | 0.72 | 1 |
| Sex-by-CBCL Externalizing | -0.011 | 0.085 | -0.13 | 0.899 | 0.91 | 0.989 | 0.84 | 1.17 |
| Sex-by-CBCL Internalizing | -0.033 | 0.078 | -0.42 | 0.676 | 0.789 | 0.968 | 0.83 | 1.13 |
| **Diffusion fMRI** |  |  |  |  |  |  |  |  |
| Intercept | 0.171 | 1.06 | 0.16 | 0.872 | 0.905 | 1.19 | 0.15 | 9.52 |
| Age | 0.009 | 0.006 | 1.53 | 0.125 | 0.309 | 1.01 | 0.99 | 1.02 |
| Male | -0.088 | 0.091 | -0.96 | 0.337 | 0.601 | 0.916 | 0.77 | 1.1 |
| Black | -0.024 | 0.143 | -0.17 | 0.869 | 0.905 | 0.976 | 0.74 | 1.29 |
| Asian | 0.154 | 0.131 | 1.17 | 0.241 | 0.482 | 1.17 | 0.90 | 1.51 |
| Other | -0.243 | 0.27 | -0.90 | 0.368 | 0.63 | 0.784 | 0.46 | 1.33 |
| Hispanic | -0.058 | 0.136 | -0.43 | 0.669 | 0.789 | 0.944 | 0.72 | 1.23 |
| Cognition | 0.004 | 0.003 | 1.41 | 0.158 | 0.359 | 1 | 0.99 | 1.01 |
| Attention | -0.05 | 0.083 | -0.61 | 0.545 | 0.738 | 0.951 | 0.81 | 1.12 |
| Externalizing | 0.047 | 0.087 | 0.54 | 0.592 | 0.756 | 1.05 | 0.88 | 1.24 |
| Internalizing | -0.011 | 0.080 | -0.14 | 0.89 | 0.91 | 0.989 | 0.85 | 1.16 |
| Sex-by-CBCL Attention | -0.334 | 0.116 | -2.89 | 0.00388 | 0.0214 | 0.716 | 0.57 | 0.89 |
| Sex-by-CBCL Externalizing | 0.051 | 0.117 | 0.44 | 0.66 | 0.789 | 1.05 | 0.84 | 1.32 |
| Sex-by-CBCL Internalizing | 0.112 | 0.108 | 1.04 | 0.299 | 0.556 | 1.12 | 0.91 | 1.38 |
| **SST task** |  |  |  |  |  |  |  |  |
| Intercept | -1.85 | 0.463 | -4 | 6.25e-05 | 0.000477 | 0.157 | 0.06 | 0.39 |
| Age | 0.014 | 0.004 | 3.92 | 8.98e-05 | 0.000629 | 1.01 | 1.01 | 1.02 |
| Male | -0.034 | 0.055 | -0.62 | 0.538 | 0.738 | 0.967 | 0.87 | 1.08 |
| Black | -0.244 | 0.085 | -2.87 | 0.00408 | 0.0214 | 0.783 | 0.66 | 0.93 |
| Asian | -0.038 | 0.081 | -0.47 | 0.636 | 0.785 | 0.963 | 0.82 | 1.13 |
| Other | -0.097 | 0.196 | -0.49 | 0.621 | 0.778 | 0.908 | 0.62 | 1.33 |
| Hispanic | -0.217 | 0.089 | -2.45 | 0.0141 | 0.0659 | 0.805 | 0.68 | 0.96 |
| Cognition | 0.016 | 0.002 | 9.2 | 3.49e-20 | 9.76e-19 | 1.02 | 1.01 | 1.02 |
| Attention | -0.139 | 0.050 | -2.77 | 0.00554 | 0.0274 | 0.87 | 0.78 | 0.96 |
| Externalizing | -0.085 | 0.053 | -1.61 | 0.108 | 0.283 | 0.919 | 0.83 | 1.02 |
| Internalizing | 0.094 | 0.048 | 1.94 | 0.0519 | 0.165 | 1.1 | 0.99 | 1.21 |
| Sex-by-CBCL Attention | -0.055 | 0.070 | -0.78 | 0.434 | 0.688 | 0.946 | 0.83 | 1.09 |
| Sex-by-CBCL Externalizing | 0.017 | 0.072 | 0.23 | 0.815 | 0.867 | 1.02 | 0.88 | 1.17 |
| Sex-by-CBCL Internalizing | -0.002 | 0.066 | -0.02 | 0.981 | 0.981 | 0.998 | 0.87 | 1.14 |
| **MID task** |  |  |  |  |  |  |  |  |
| Intercept | 0.758 | 0.51 | 1.49 | 0.137 | 0.322 | 2.13 | 0.78 | 5.8 |
| Age | 0.004 | 0.004 | 0.90 | 0.366 | 0.63 | 1 | 0.99 | 1.01 |
| Male | -0.25 | 0.061 | -4.09 | 4.31e-05 | 0.000362 | 0.779 | 0.69 | 0.88 |
| Black | -0.163 | 0.095 | -1.72 | 0.0863 | 0.242 | 0.85 | 0.71 | 1.02 |
| Asian | 0.189 | 0.092 | 2.05 | 0.0407 | 0.143 | 1.21 | 1.01 | 1.45 |
| Other | -0.313 | 0.198 | -1.58 | 0.114 | 0.289 | 0.731 | 0.49 | 1.08 |
| Hispanic | -0.053 | 0.099 | -0.53 | 0.594 | 0.756 | 0.948 | 0.78 | 1.15 |
| Cognition | 0.007 | 0.002 | 3.67 | 0.000244 | 0.00158 | 1.01 | 1 | 1.01 |
| Attention | -0.064 | 0.058 | -1.12 | 0.262 | 0.512 | 0.938 | 0.84 | 1.05 |
| Externalizing | 0.019 | 0.061 | 0.31 | 0.758 | 0.849 | 1.02 | 0.91 | 1.15 |
| Internalizing | -0.015 | 0.056 | -0.27 | 0.791 | 0.861 | 0.985 | 0.88 | 1.1 |
| Sex-by-CBCL Attention | -0.056 | 0.078 | -0.72 | 0.473 | 0.698 | 0.946 | 0.81 | 1.1 |
| Sex-by-CBCL Externalizing | -0.051 | 0.081 | -0.64 | 0.524 | 0.734 | 0.95 | 0.81 | 1.11 |
| Sex-by-CBCL Internalizing | 0.058 | 0.074 | 0.78 | 0.435 | 0.688 | 1.06 | 0.92 | 1.22 |
| **EN-Back task** |  |  |  |  |  |  |  |  |
| Intercept | -5.16 | 0.467 | -11.1 | 2.14e-28 | 9e-27 | 0.006 | 0.002 | 0.01 |
| Age | 0.027 | 0.004 | 7.65 | 1.95e-14 | 4.1e-13 | 1.03 | 1.02 | 1.04 |
| Male | 0.045 | 0.053 | 0.84 | 0.4 | 0.673 | 1.05 | 0.94 | 1.16 |
| Black | -0.348 | 0.082 | -4.23 | 2.33e-05 | 0.000245 | 0.706 | 0.60 | 0.83 |
| Asian | -0.17 | 0.078 | -2.19 | 0.0286 | 0.114 | 0.844 | 0.72 | 0.98 |
| Other | -0.199 | 0.194 | -1.03 | 0.305 | 0.556 | 0.82 | 0.56 | 1.2 |
| Hispanic | -0.157 | 0.089 | -1.77 | 0.0769 | 0.223 | 0.855 | 0.72 | 1.02 |
| Cognition | 0.032 | 0.002 | 17.2 | 2.23e-66 | 1.87e-64 | 1.03 | 1.03 | 1.04 |
| Attention | -0.142 | 0.049 | -2.93 | 0.0034 | 0.0204 | 0.868 | 0.78 | 0.95 |
| Externalizing | -0.115 | 0.051 | -2.24 | 0.0253 | 0.112 | 0.891 | 0.81 | 0.99 |
| Internalizing | 0.055 | 0.047 | 1.18 | 0.239 | 0.482 | 1.06 | 0.97 | 1.16 |
| Sex-by-CBCL Attention | 0.075 | 0.068 | 1.1 | 0.273 | 0.522 | 1.08 | 0.94 | 1.23 |
| Sex-by-CBCL Externalizing | 0.029 | 0.071 | 0.41 | 0.686 | 0.789 | 1.03 | 0.89 | 1.18 |
| Sex-by-CBCL Internalizing | 0.017 | 0.065 | 0.26 | 0.798 | 0.861 | 1.02 | 0.89 | 1.15 |

Note: CBCL, Child Behavior Checklist; EN-Back, Emotional N-Back Task; MID, Monetary Incentive Delay Task; OR, odds ratio; SST, Stop Signal Task; General Cognition is measured by the NIH Toolbox age corrected scores(3). Odds ratios are reported relative to the reference group. Odds ratios are reported relative to the reference group in the case of categorical variables. Odds ratios > 1 indicate increased likelihood of passing quality control among the nonreference group relative to the reference group (i.e., lower likelihood of passing quality control in the reference group). While odds ratios < 1 indicate decreased likelihood of passing quality control. For continuous measures, the odds ratios represent the change in odds per unit change in the measure.

**Table S3:** Results of Linear Mixed-Effects Models Predicting Motion and Domains of Transdiagnostic Symptoms with Sex Interactions

| **Variables** | **Estimate** | **Std. Error** | **Cohen's D** | ***t* value** | ***p*** | ***p*_FDR_** | **95% CI** | |
| --- | --- | --- | --- | --- | --- | --- | --- | --- |
|  |  |  |  |  |  |  | **2.5 %** | **97.5 %** |
| **Resting-state fMRI** |  |  |  |  |  |  |  |  |
| Intercept | 0.824 | 0.046 | 0.533 | 17.9 | 6.29e-69 | 4.41e-67 | 0.733 | 0.914 |
| Age | -0.003 | 0.0004 | -0.215 | -9.45 | 4.56e-21 | 4.56e-20 | -0.004 | -0.003 |
| Male | 0.052 | 0.009 | 0.137 | 5.61 | 2.07e-08 | 8.05e-08 | 0.034 | 0.071 |
| Black | 0.051 | 0.008 | 0.165 | 6.22 | 5.42e-10 | 2.37e-09 | 0.035 | 0.067 |
| Asian | 0.021 | 0.009 | 0.058 | 2.32 | 0.0206 | 0.0379 | 0.003 | 0.040 |
| Other | -0.001 | 0.0002 | -0.181 | -7.81 | 6.75e-15 | 3.94e-14 | -0.002 | -0.001 |
| Hispanic | 0.029 | 0.020 | 0.034 | 1.47 | 0.143 | 0.208 | -0.010 | 0.068 |
| Cognition | 0.028 | 0.005 | 0.119 | 5.17 | 2.42e-07 | 8.47e-07 | 0.018 | 0.039 |
| Attention | 0.015 | 0.005 | 0.070 | 3.06 | 0.00225 | 0.00543 | 0.005 | 0.024 |
| Externalizing | 0.013 | 0.005 | 0.056 | 2.44 | 0.0148 | 0.0305 | 0.002 | 0.023 |
| Internalizing | -0.017 | 0.005 | -0.079 | -3.46 | 0.000537 | 0.00145 | -0.026 | -0.007 |
| Sex-by-CBCL Attention | 0.004 | 0.007 | 0.012 | 0.52 | 0.606 | 0.663 | 0.101 | 0.197 |
| Sex-by-CBCL Externalizing | -0.002 | 0.007 | -0.007 | -0.31 | 0.753 | 0.787 | 0.129 | 0.254 |
| Sex-by-CBCL Internalizing | 0.004 | 0.007 | 0.015 | 0.63 | 0.526 | 0.604 | 0.401 | 0.436 |
| **Diffusion fMRI** |  |  |  |  |  |  |  |  |
| Intercept | 1.73 | 0.098 | 1.65 | 17.7 | 4.06e-54 | 9.48e-53 | 1.54 | 1.92 |
| Age | -0.002 | 0.0007 | -0.066 | -2.82 | 0.00478 | 0.0105 | -0.003 | -0.0006 |
| Male | 0.061 | 0.018 | 0.087 | 3.42 | 0.000639 | 0.00166 | 0.026 | 0.097 |
| Black | 0.046 | 0.016 | 0.072 | 2.85 | 0.00437 | 0.00986 | 0.014 | 0.078 |
| Asian | 0.030 | 0.018 | 0.043 | 1.65 | 0.0982 | 0.164 | -0.006 | 0.066 |
| Other | -0.002 | 0.0003 | -0.127 | -5.37 | 7.91e-08 | 2.91e-07 | -0.002 | -0.001 |
| Hispanic | 0.062 | 0.040 | 0.038 | 1.56 | 0.12 | 0.186 | -0.016 | 0.139 |
| Cognition | 0.008 | 0.011 | 0.019 | 0.79 | 0.43 | 0.518 | -0.013 | 0.029 |
| Attention | 0.014 | 0.010 | 0.035 | 1.49 | 0.136 | 0.203 | -0.019 | 0.034 |
| Externalizing | 0.006 | 0.010 | 0.014 | 0.62 | 0.536 | 0.606 | -0.035 | 0.021 |
| Internalizing | -0.015 | 0.009 | -0.038 | -1.61 | 0.107 | 0.175 | -0.016 | 0.035 |
| Sex-by-CBCL Attention | 0.007 | 0.013 | 0.013 | 0.55 | 0.582 | 0.646 | 0.080 | 0.164 |
| Sex-by-CBCL Externalizing | -0.007 | 0.014 | -0.011 | -0.49 | 0.625 | 0.673 | 0.049 | 0.098 |
| Sex-by-CBCL Internalizing | 0.009 | 0.013 | 0.017 | 0.724 | 0.469 | 0.55 | 0.349 | 0.379 |
| **SST task** |  |  |  |  |  |  |  |  |
| Intercept | 1.09 | 0.079 | 0.411 | 13.8 | 2.19e-42 | 3.07e-41 | 0.938 | 1.25 |
| Age | -0.004 | 0.0006 | -0.155 | -6.52 | 7.43e-11 | 3.47e-10 | -0.005 | -0.003 |
| Male | 0.051 | 0.017 | 0.078 | 3.09 | 0.00203 | 0.00509 | 0.019 | 0.083 |
| Black | 0.055 | 0.014 | 0.108 | 3.86 | 0.000114 | 0.000319 | 0.027 | 0.083 |
| Asian | 0.039 | 0.016 | 0.062 | 2.36 | 0.0183 | 0.0347 | -0.005 | 0.029 |
| Other | -0.002 | 0.0003 | -0.197 | -8.18 | 3.37e-16 | 2.14e-15 | 0.001 | 0.037 |
| Hispanic | 0.007 | 0.033 | 0.005 | 0.22 | 0.823 | 0.835 | -0.057 | 0.072 |
| Cognition | 0.054 | 0.009 | 0.137 | 5.7 | 1.23e-08 | 5.08e-08 | -0.025 | 0.008 |
| Attention | 0.012 | 0.008 | 0.034 | 1.42 | 0.155 | 0.222 | -0.011 | 0.036 |
| Externalizing | 0.019 | 0.009 | 0.050 | 2.11 | 0.0348 | 0.0609 | -0.040 | 0.009 |
| Internalizing | -0.008 | 0.008 | -0.024 | -1.01 | 0.312 | 0.398 | -0.020 | 0.025 |
| Sex-by-CBCL Attention | 0.012 | 0.012 | 0.025 | 1.05 | 0.295 | 0.383 | 0.032 | 0.13 |
| Sex-by-CBCL Externalizing | -0.016 | 0.013 | -0.030 | -1.27 | 0.204 | 0.28 | 0.043 | 0.084 |
| Sex-by-CBCL Internalizing | 0.003 | 0.012 | 0.006 | 0.23 | 0.816 | 0.835 | 0.325 | 0.35 |
| **MID task** |  |  |  |  |  |  |  |  |
| Intercept | 1.16 | 0.069 | 0.486 | 16.7 | 3.36e-61 | 1.18e-59 | 1.03 | 1.3 |
| Age | -0.005 | 0.0005 | -0.2 | -8.72 | 3.52e-18 | 2.74e-17 | -0.006 | -0.004 |
| Male | 0.069 | 0.014 | 0.121 | 4.88 | 1.07e-06 | 3.25e-06 | -0.003 | 0.052 |
| Black | 0.061 | 0.012 | 0.135 | 4.92 | 9.08e-07 | 2.89e-06 | -0.003 | -0.002 |
| Asian | 0.025 | 0.014 | 0.045 | 1.75 | 0.0806 | 0.138 | 0.006 | 0.035 |
| Other | -0.003 | 0.0003 | -0.229 | -9.83 | 1.12e-22 | 1.31e-21 | -0.003 | 0.028 |
| Hispanic | -0.022 | 0.030 | -0.017 | -0.72 | 0.471 | 0.55 | 0.055 | 0.087 |
| Cognition | 0.071 | 0.008 | 0.199 | 8.56 | 1.32e-17 | 9.26e-17 | -0.022 | 0.006 |
| Attention | 0.020 | 0.007 | 0.063 | 2.73 | 0.00629 | 0.0134 | -0.005 | 0.036 |
| Externalizing | 0.013 | 0.008 | 0.037 | 1.6 | 0.11 | 0.175 | -0.031 | 0.012 |
| Internalizing | -0.008 | 0.007 | -0.026 | -1.11 | 0.265 | 0.35 | -0.020 | 0.019 |
| Sex-by-CBCL Attention | 0.016 | 0.010 | 0.034 | 1.5 | 0.134 | 0.203 | 0.141 | 0.208 |
| Sex-by-CBCL Externalizing | -0.009 | 0.011 | -0.019 | -0.83 | 0.407 | 0.509 | 0.048 | 0.094 |
| Sex-by-CBCL Internalizing | -0.0006 | 0.010 | -0.001 | -0.06 | 0.954 | 0.954 | 0.328 | 0.362 |
| **EN-Back task** |  |  |  |  |  |  |  |  |
| Intercept | 1.19 | 0.081 | 0.431 | 14.6 | 1.74e-47 | 3.05e-46 | 1.03 | 1.35 |
| Age | -0.005 | 0.0006 | -0.177 | -7.23 | 5.5e-13 | 2.75e-12 | -0.006 | -0.003 |
| Male | 0.072 | 0.017 | 0.108 | 4.19 | 2.78e-05 | 8.11e-05 | 0.039 | 0.106 |
| Black | 0.073 | 0.015 | 0.145 | 4.99 | 6.25e-07 | 2.08e-06 | 0.044 | 0.102 |
| Asian | 0.050 | 0.017 | 0.081 | 3.03 | 0.00247 | 0.00577 | 0.018 | 0.083 |
| Other | -0.003 | 0.0003 | -0.216 | -8.85 | 1.1e-18 | 9.6e-18 | -0.003 | -0.002 |
| Hispanic | 0.046 | 0.034 | 0.035 | 1.38 | 0.168 | 0.235 | -0.020 | 0.112 |
| Cognition | 0.075 | 0.010 | 0.188 | 7.71 | 1.4e-14 | 7.53e-14 | 0.056 | 0.094 |
| Attention | 0.021 | 0.009 | 0.059 | 2.41 | 0.0159 | 0.0309 | 0.004 | 0.038 |
| Externalizing | 0.021 | 0.009 | 0.055 | 2.27 | 0.023 | 0.0412 | 0.003 | 0.040 |
| Internalizing | -0.021 | 0.009 | -0.059 | -2.43 | 0.0152 | 0.0305 | -0.038 | -0.004 |
| Sex-by-CBCL Attention | 0.015 | 0.012 | 0.029 | 1.19 | 0.234 | 0.315 | -0.009 | 0.038 |
| Sex-by-CBCL Externalizing | -0.010 | 0.013 | -0.020 | -0.81 | 0.418 | 0.513 | -0.036 | 0.015 |
| Sex-by-CBCL Internalizing | 0.004 | 0.012 | 0.009 | 0.37 | 0.713 | 0.756 | -0.019 | 0.028 |

Note: CBCL, Child Behavior Checklist; EN-Back, Emotional N-Back Tas; MID, Monetary Incentive Delay Task; SST, Stop Signal Task. General Cognition is measured by the NIH Toolbox age corrected scores(3).

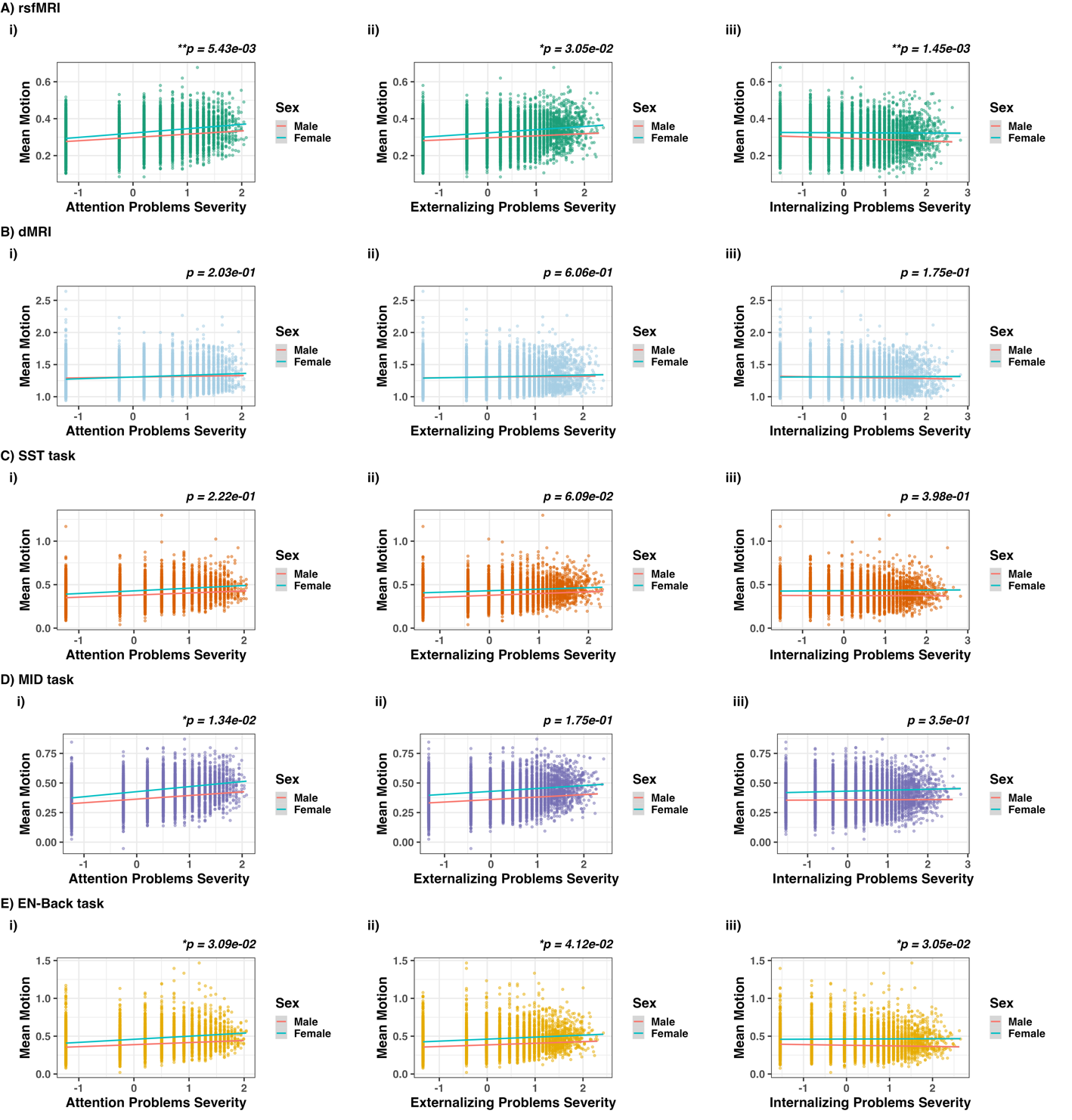
**Figure S1.** Sex differences in severity of transdiagnostic symptom domains in youths were not associated with motion during functional and structural imaging. Scatterplots depict results of linear mixed-effects models (see Table 3) for CBCL Attention, Externalizing, and Internalizing Problem scores for resting-state fMRI (A), diffusion MRI (B), Stop Signal Task (SST) (C), Monetary Incentive Delay (MID) task (D), and the emotional version of the n-back task (EN-Back) (E). The red trendline line represents the regression line based on the linear mixed-effects model fit. Statistical significance is denoted by asterisks: ***p < 0.001, **p < 0.01, and *p < 0.05.

**Supplement Discussion:**

**Associations with Demographic Characteristics**

Our findings showed that younger age and reduced cognitive performance were associated with greater levels of motion. These findings related to age are consistent with prior work indicating inverse relationships between age and motion in pediatric samples (4-10). However, other studies have reported a positive relationship between age and motion in child and adult samples (11, 12). We also found greater motion in males relative to females, which aligns with prior studies (4, 10, 11, 13, 14). However, studies have also reported mixed and/or null findings related to sex effects and motion (5, 6). It is important to note that our sample was relatively gender-balanced (51% males; 49% females) and consisted of a large-scale pediatric sample (>9,000 youths) with varying levels of symptom severities (1). Additionally, it has also been suggested that demographic groups may be an unstable predictor of motion due to inter-individual variance in motion (4). Thus, these findings should be interpreted with caution and are provided to facilitate comparison with prior work.
